# Kinesin-induced buckling reveals the limits of microtubule self-repair

**DOI:** 10.1101/2025.09.08.672697

**Authors:** Shweta Nandakumar, Jonas Bosche, Mirko Wieczorek, Constantin Matteo Albrecht, Belinda König, Mona Grünewald, Ludger Santen, Stefan Diez, Reza Shaebani, Laura Schaedel

## Abstract

Microtubules are stiff cytoskeletal polymers whose ability to rapidly switch between growth and disassembly relies on a metastable lattice. This metastability is also reflected in their sensitivity to environmental conditions and in intrinsic lattice dynamics, where spontaneous tubulin loss is balanced by tubulin incorporation from solution – a process that also enables microtubules to self-repair when damaged. Whether such intrinsic self-repair is sufficient to preserve microtubule integrity during dynamic molecular-motor induced buckling, which frequently occurs in cells, remains unclear. Here, we show that kinesin-driven microtubule buckling *in vitro* induces severe lattice damage, leading to extensive tubulin incorporation. In many cases, however, the damage exceeds the microtubules’ capacity for self-repair, resulting in breakage. In contrast, microtubules survive continuous buckling substantially longer in the presence of intracellular factors. Our results identify the limits of intrinsic microtubule self-repair and demonstrate that additional cellular mechanisms are essential to maintain microtubule integrity under sustained mechanical load.

## Introduction

Microtubules are highly dynamic cytoskeletal filaments that support intracellular organization and adapt rapidly to physiological cues [1]. Microtubule tip dynamics – alternating between growth and disassembly – is driven by GTP hydrolysis [1,2], which renders the microtubule lattice inherently metastable and particularly sensitive to environmental conditions such as temperature [3,4]. This inherent fragility not only enables microtubules to remain responsive to signals, but also makes them vulnerable to destabilization.

A manifestation of this metastability is intrinsic lattice dynamics, wherein tubulin subunits dissociate from the lattice, particularly at structural defects, and are replaced by free tubulin from solution [5]. This process occurs slowly but can be accelerated by repeated bending via orthogonal fluid flow [6], the activity of severing enzymes and the non-enzymatic microtubule-associated protein (MAP) tau [7,8], and even the translocation of unloaded motor proteins [9–11]. Although other MAPs such as CLASP[12] and CLIP-170 [13] may support repair or stabilization, it is generally assumed that microtubules have sufficient intrinsic self-repair capacity to withstand damage [6,14,15]. However, in cells, microtubules are frequently exposed to strong and dynamic deformations [16,17], including pronounced buckling caused by opposing motor forces and anchorage points within the cytoplasm [18–24]. Whether intrinsic self-repair is sufficient to maintain microtubule integrity under such sustained mechanical stress is unclear.

Here, we use a combination of *in vitro* reconstitution, a stochastic computational model, and supporting cellular data to investigate microtubule behavior under motor-induced buckling. We find that buckling leads to accelerated lattice damage and extensive tubulin incorporation, and that damage frequently exceeds the capacity of self-repair, resulting in microtubule breakage. However, in the presence of intracellular factors, microtubules are significantly more resilient. These findings reveal the limits of intrinsic microtubule self-repair and highlight the essential contribution of cellular mechanisms in preserving microtubule integrity under persistent mechanical load.

## Results

### 1) Static curvature induces microtubule self-repair

Despite their high flexural rigidity [25], microtubules frequently adopt curved conformations in cells [26–28], suggesting that they are exposed to considerable intracellular forces. For example, in fixed PtK2 cells with endogenously labeled tubulin, we observed that many microtubules display locally highly curved regions (**Fig. 1a**). This raises the question whether static bending promotes lattice damage and repair. To test this, we reconstituted statically curved microtubules *in vitro* (**Fig. 1b**): First, we polymerized dynamic microtubules from stabilized biotinylated seeds (step I) and stabilized their ends with biotin-tubulin caps (step II). Importantly, the GDP microtubule lattice between seed and cap was not stabilized. These microtubules were then introduced into streptavidin-coated flow chambers assembled from passivated cover-glass (step III). We alternated the flow direction during chamber loading, frequently resulting in microtubules adopting bent conformations upon immobilization at both ends (step IV). The resulting microtubule curvatures are comparable to that of intracellular microtubules (**ED Fig. 1d**). Next, we exposed these microtubules to soluble fluorescently labeled tubulin for 15 min to allow for incorporation (step V), followed by washout and imaging (step VI). We observed tubulin incorporation both in straight and bent microtubules, where incorporation events appear as localized stretches along the microtubule lattice (**Fig. 1c, ED Fig. 1a**). While the length of individual incorporation stretches does not differ between bent and straight microtubules when analyzed along the entire microtubule (**ED Fig. 1b**), focusing specifically on bent zones reveals a modest increase in incorporation length compared to straight microtubules (**Fig. 1d(i)**). In contrast, the spatial frequency of incorporation events is substantially higher in bent microtubules (**Fig. 1d(ii)**), resulting in 28% of lattice lengths showing tubulin incorporation, compared to only 4% in straight microtubules (**Fig. 1d(iii)**). When grouping microtubules by curvature, we found a progressive increase in tubulin incorporation with increasing curvature (**Fig. 1e**). These observations suggest that static bending promotes lattice damage and subsequent self-repair *in vitro*.

**Figure 1:**
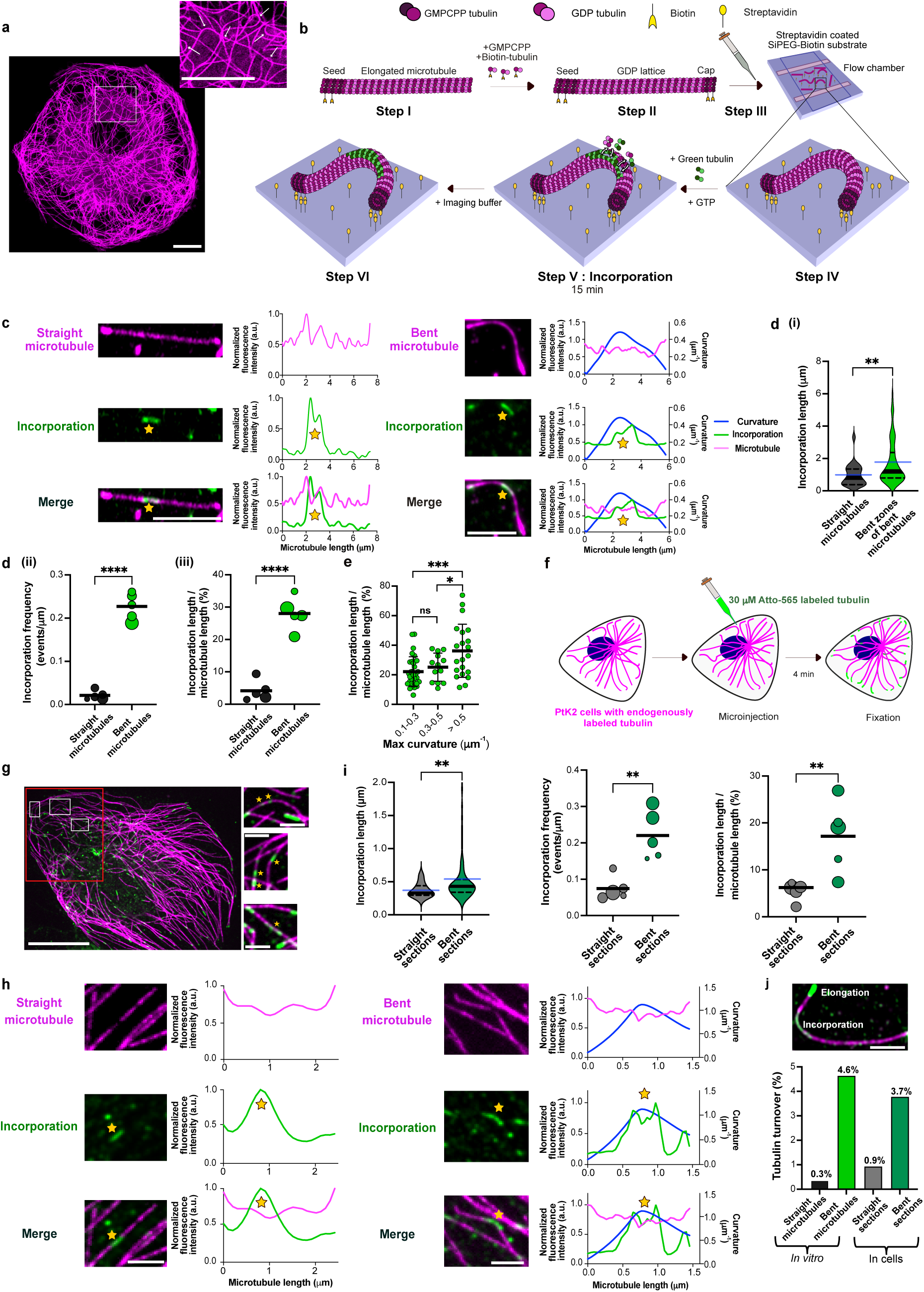
Static curvature triggers microtubule damage and consequent self-repair. **a**, Microtubules adopt bent and buckled conformations in cells: Image of bent microtubules in a PtK2 cell (tubulin-eGFP, represented in magenta). Inset shows a zoomed-in image with white arrows indicating bent microtubules. Scale bars: 10 μm. **b**, Schematic of the *in vitro* experimental setup used to assess self-repair in static bent microtubules. Capped GDP microtubules (shown in magenta) with biotinylated GMPCPP seeds and caps (Step I and II) were flushed onto a streptavidin coated flow chamber made from SiPEG-biotin passivated cover-glasses (Step III). Microtubules of various curvatures were obtained by alternating the flow direction (Step IV). Microtubules were then incubated with 5 μM green labeled tubulin for 15 min (Step V) followed by washing with imaging buffer (Step VI). **c**, Example images showing incorporation of green labeled tubulin (marked with a yellow star) in a straight and in a bent microtubule *in vitro*. Scale bar: 5 μm. Graphs represent line scans of the microtubule (magenta), the incorporation channel (green) and curvature (blue). Profiles have been normalized to 1 for the maximum value of the intensity (a.u.) for the microtubule and incorporation channel, respectively **d**, **(i):** Violin plot showing longer incorporation stretches in zones of high local curvature in bent microtubules *in vitro*. Total length of microtubules analyzed: 1,104 μm from three independent experiments (p= 0.0069 using Mann-Whitney test) (n= 23 incorporations for straight microtubules, n= 83 incorporations for bent microtubules). Black solid line represents the median and dotted lines represent the interquartile range. Blue line represents the mean. **(ii):** Bubble plot showing higher frequency of incorporations in bent microtubules when compared to straight microtubules *in vitro*. Bubble sizes scale with the total microtubule length analyzed. Each circle represents an independent dataset (comprising of 291, 295, 350, 165 and 65 μm of total microtubule length analyzed for bent microtubules and 311, 347, 340, 230 and 366 μm of total microtubule length analyzed for straight microtubules). The black line represents the mean. p<0.0001 using unpaired t-test. **(iii):** Bubble plot showing higher amount of lattice turnover, estimated as incorporation length/ microtubule length, in bent microtubules when compared to straight microtubules *in vitro*. Bubble sizes scale with the total microtubule length analyzed. Each circle represents an independent dataset (comprising 291, 295, 350, 165 and 65 μm of the total microtubule length analyzed for bent microtubules and 311, 347, 340, 230 and 366 μm of the total microtubule length analyzed for straight microtubules). The black line represents the mean. p<0.0001 using unpaired t-test. **e**, Scatter dot plot comparing the lattice length with incorporation in bent microtubules across different curvature ranges *in vitro*. Total length of microtubules analyzed: 1,104 μm from three independent experiments. Black lines represent the mean and S.D. p= 0.3597 (not significant; for curvatures 0.1-0.3 μm^-1^ and 0.3-0.5 μm^-1^), p= 0.0006 (for curvatures > 0.5 μm^-1^ and 0.1-0.3 μm^-1^) and p= 0.0493 (for curvatures > 0.5 μm^-1^ and 0.3-0.5 μm^-1^) using unpaired t-test. **f,** Schematic of experimental setup used by Gazzola et al., 2023 to assess microtubule self-repair in PtK2 cells. 30 μM of ATTO-565 labeled tubulin (represented here in green) was microinjected into PtK2 cells expressing endogenous tubulin-eGFP (represented here in magenta) and the cells were fixed after 4 min and imaged. **g,** Example images showing microtubule self-repair in a PtK2 cell, scale bar: 20 μm. Box with red outline indicates a selected region of interest. Insets show zoomed-in images of microtubule sections with incorporations marked with a yellow star. Scale bars of inset images: 2 μm. **h**, Self-repair (marked with a yellow star) in both straight and bent microtubule sections in cells. Scale bar: 2 μm. Graphs represent line scans of the microtubule (magenta), curvature (blue) and the incorporation channel (green). Profiles have been normalized to 1 for the maximum value of the intensity (a.u.) for the microtubule and incorporation channel. **i**, **Left:** Violin plot showing longer incorporation stretches in bent microtubule sections in cells. Total length of microtubules analyzed: 401.96 μm from five cells (p= 0.0025 using Mann-Whitney test.) (n= 37 incorporations for straight sections and n= 84 incorporations for bent sections). Black line represents the median and dotted lines represent the interquartile range. Blue line represents the mean. **Center:** Bubble plot showing higher frequency of incorporations in bent microtubule sections when compared to straight microtubule sections in cells. Bubble sizes scale with the total microtubule length analyzed. Each circle represents an independent dataset from one cell (comprising of 130, 98, 57, 44 and 72 μm of total length analyzed for bent sections and 22, 56, 74, 36 and 153 μm of total microtubule length analyzed for straight sections. Black line represents the mean. p= 0.0022 using unpaired t-test. **Right:** Bubble plot showing higher amount of lattice length with incorporation estimated as incorporation length/microtubule length, in bent microtubule sections when compared to straight microtubule sections in cells. Bubble sizes scale with the total microtubule length analyzed. Each circle represents an independent dataset from one cell (comprising of 130, 98, 57, 44 and 72 μm of total length analyzed for bent sections and 22, 56, 74, 36 and 153 μm of total microtubule length analyzed for straight sections). Black lines represent the mean. p= 0.0099 using unpaired t-test. **j**, **Top:** Example image showing an incorporation stretch and the elongated tip that was used as a reference to estimate the amount of lateral tubulin incorporation (see Methods). Scale bar: 2 μm. **Bottom**: Higher % of tubulin turnover in bent microtubules, both in cells and *in vitro*. Three independent experiments were analyzed for each condition. n= 21 incorporations for straight microtubules (*in vitro)*, n= 51 incorporations for bent microtubules (*in vitro)*, n= 16 incorporations for straight sections (cells) and n= 32 incorporations for bent sections (cells). Refer methods for estimation of % tubulin turnover and **ED Fig. 1e** for estimation of amount of lateral tubulin incorporation.

We then proceeded to assess the relationship between curvature and tubulin incorporation in cells by reanalyzing a previously published dataset [29] of self-repair in PtK2 cells expressing endogenously GFP-tagged tubulin (represented here in magenta). These cells were micro-injected with 30 µM purified, Atto-565 labeled tubulin (represented here in green), followed by fixation after 4 min of incubation (**Fig. 1f,g**). Since globally straight microtubules are rare in cells, we decided to analyze microtubules section-wise (see methods). Similar to our observations *in vitro*, we detected incorporation events along both straight and bent microtubule sections (**Fig. 1h, ED Fig. 1c**). Tubulin incorporation stretches are longer (**Fig. 1i, left**) and more frequent (**Fig. 1i, center**) in bent microtubule sections compared to straight microtubule sections, overall leading to a larger proportion of lattice lengths with incorporations (**Fig. 1i, right**), reminiscent of our *in vitro* results.

Since incorporation stretches appear more intense in intracellular microtubules compared to *in vitro* microtubules (suggesting a higher fraction of the lattice has been replaced at these sites), we then estimated the local amount of incorporated tubulin across protofilaments *in vitro* and in cells (see methods). For this, we normalized the intensities of the incorporation stretches to the intensities of microtubule tips grown with tubulin of the same color, which we presume to consist of 13 protofilaments (**Fig. 1j, top**). This quantification serves as a direct readout of the lateral extent of tubulin incorporation, revealing that incorporation typically occurs over 1-3 protofilaments, occasionally reaching up to 9 protofilaments in cells (**ED Fig. 1e**). Thus, in most cases, only a small portion of the lattice is exchanged. Together with the mean length of lattice showing incorporation (**Fig. 1d(iii) and Fig. 1i, right**), this analysis allowed us to estimate the total amount of tubulin turnover in each condition by accounting for both the longitudinal extent of incorporation along the microtubule and its lateral spread across protofilaments. **Fig. 1j (bottom)** shows that tubulin turnover is more pronounced in bent as opposed to straight microtubules, both *in vitro* and in cells, yet remains below 5%. Together, these data show that microtubule curvature leads to increased tubulin incorporation both *in vitro* and in cells. While previous work has shown that repeated bending cycles by orthogonal fluid flow can induce microtubule damage [6], our findings demonstrate that sustained static curvature is sufficient to enhance tubulin incorporation, indicating a direct mechanical contribution to lattice damage and repair.

### 2) *In vitro* reconstitution of cell-like microtubule buckling

While the striking similarity between tubulin incorporation *in vitro* and in cells highlights the role of microtubule curvature, our *in vitro* observations were made after an incubation time three times longer than that used in cells (15 min vs. 4 min). This discrepancy suggests that additional factors contribute to microtubule damage and repair in cells. When examining dynamic microtubules in live cells, it becomes apparent that local microtubule curvature often changes over time (**Fig. 2a; suppl. movie 1**), as has been described earlier [18]. While some microtubules maintain a curved conformation with little change over extended periods (**Fig. 2b (i)**), many exhibit dynamic buckling on short timescales (**Fig. 2b (ii)**). In rare instances, we observed microtubule breakage, typically occurring in regions of high curvature (**Fig. 2b (iii), ED Fig. 3b**). Refer **ED Fig. 3a** for quantification of different microtubule bending events in cells. This prompted us to investigate how dynamic microtubule buckling affects microtubule integrity.

**Figure 2:**
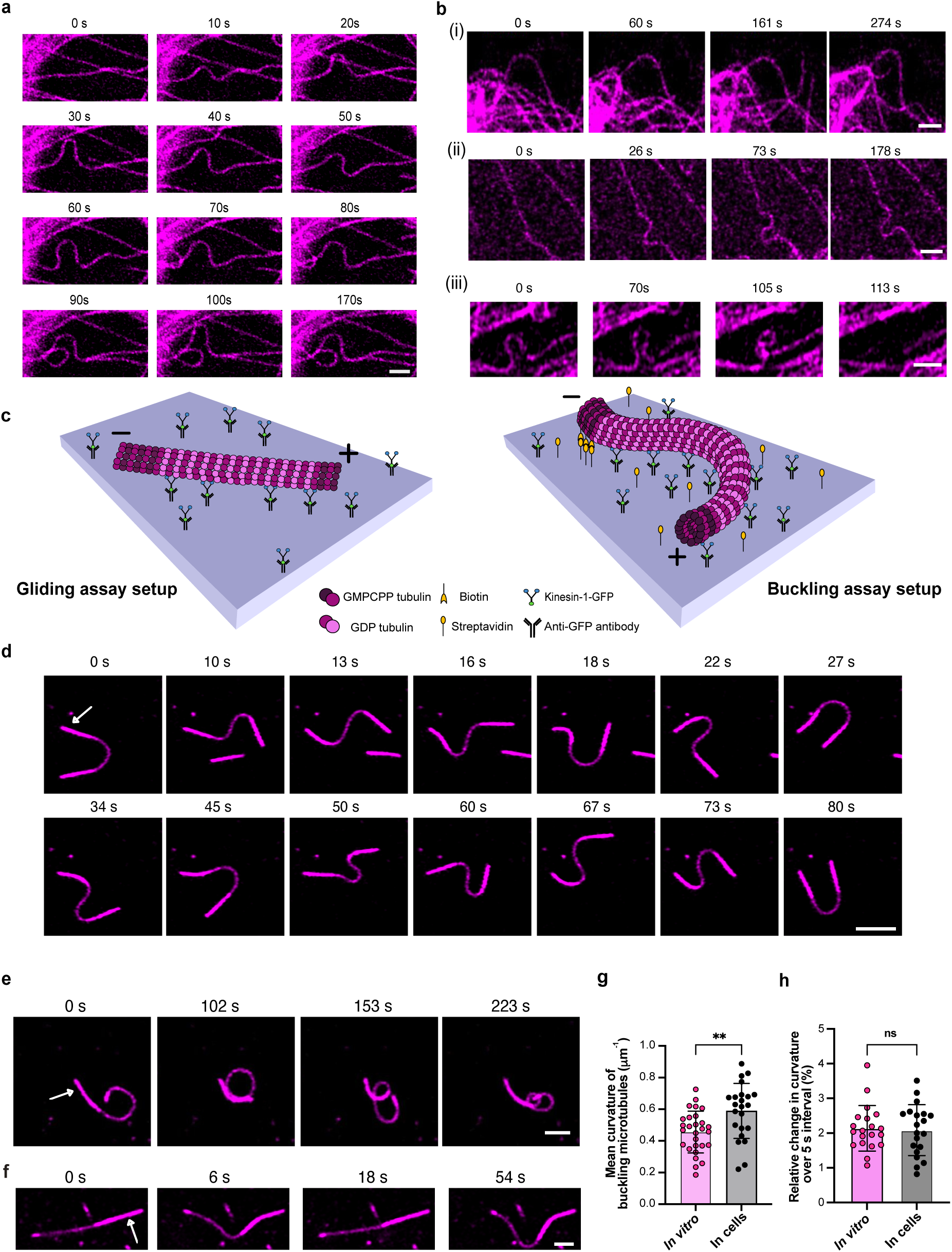
*In vitro* reconstitution of cell-like microtubule buckling. **a**, Time-lapse sequence showing change in local microtubule curvature over time in endogenously labeled PtK2 cells (tubulin-eGFP, represented in magenta). Scale bar: 2 μm. **b**, Time-lapse sequence of microtubule bending and buckling events in PtK2 cells: **(i)** curved microtubule persisting in a bent form; **(ii)** dynamically buckling microtubule; **(iii)** breakage of a looping microtubule. Scale bars: 2 μm. **c**, Schematic of the experimental setup used to reconstitute buckling *in vitro*. In typical gliding assays (left), kinesin-1-GFP motor proteins are immobilized onto an anti-GFP antibody coated glass surface and GDP microtubules glide over the layer of motors in the presence of ATP. In the buckling assay setup (right), the surface is coated with equal amounts of streptavidin and anti-GFP antibody. Using capped microtubules with seeds containing biotin, we immobilized one end (minus end) to the surface. When the mix with ATP is added, microtubules buckle dynamically. **d**, Time-lapse sequence of a microtubule displaying regular flagella-like oscillations. White arrow in the first frame indicates the position of the microtubule seed (minus end). Scale bar: 5 μm. **e**, Time-lapse sequence of a looping microtubule. White arrow in the first frame indicates the position of the microtubule seed (minus end). Scale bar: 2 μm. **f**, Microtubule showing a regular beating pattern. White arrow in the first frame indicates the position of the microtubule seed (minus end). Scale bar: 2 μm. **g**, Comparison of mean curvature of buckling microtubules in cells and *in vitro* (n= 28 timepoints, *in vitro* from 4 buckling microtubules from three independent experiments and n= 23 timepoints from 4 cells analyzed from 2 independent experiments). p= 0.0032 using unpaired t-test. Error bars represent the S.D. **h**, Comparison of rate of change in curvature in buckling microtubules in 5 s in cells and *in vitro* (n= 20 from three independent experiments for *in vitro* and from 5 cells analyzed from 2 independent experiments). p= 0.8125 (not significant; ns), using unpaired t-test. Error bars represent the S.D.

Since the origin and magnitude of intracellular forces are difficult to determine, we sought to reconstitute microtubule buckling *in vitro*. Given that the dynamic, short-wavelength microtubule buckling in cells has been mainly attributed to molecular motor activity [17,23,26], we adapted the motor-based microtubule gliding assay setup to mimic this behavior (**Fig. 2c**) [30–32]. We first grew dynamic microtubules from biotinylated seeds and stabilized their ends with biotin-free GMPCPP-tubulin. We then introduced these microtubules into flow chambers coated with streptavidin and GFP-labeled kinesin-1 (immobilized via anti-GFP antibodies). Since the biotinylated seeds typically elongate at their plus ends, microtubules are effectively anchored at their minus ends to the streptavidin coated surface. When ATP is added, motor-driven translocation of the microtubule shaft induces microtubule buckling.

In our *in vitro* buckling assay, microtubules exhibit a range of dynamic deformation modes (**Fig. 2d-f; suppl. movie 2**). To describe this diversity more systematically, we distinguish three recurring deformation patterns: some microtubules show regular, flagella-like oscillations (**Fig. 2d**), characterized by a repeating motion pattern with regions of high curvature that originate near the anchor point and travel towards the free end (see **suppl. movie 3**). Others form chaotic loops without any apparent repeating pattern (**Fig. 2e**). Some also display a beating-like motion (**Fig. 2f**), which also repeats at regular intervals but lacks traveling regions of high curvature, and intermittently returns to relatively straight conformations. To compare these deformation dynamics to those observed in PtK2 cells, we measured microtubule curvatures and rates of curvature change in both systems. The distributions are similar, with slightly higher curvatures in cells, indicating that our assay reproduces the dynamic shape fluctuations of buckling microtubules seen in cells (**Fig. 2g,h**), and reaches curvatures comparable to those of the bent zones in static bent microtubules (**ED Fig. 3c**). While the precise intracellular force patterns remain elusive, these observations suggest that key aspects of motor-induced buckling can be reconstituted using a simplified *in vitro* system.

### 3) Kinesin-driven buckling causes extensive damage and self-repair

We then used our *in vitro* assay to study microtubule damage and self-repair in buckling microtubules (**Fig. 3a**). For this, we let capped GDP microtubules (magenta) buckle dynamically in the presence of free tubulin (green) for 15 min before washing out the labeled free tubulin and imaging. **Fig. 3b** shows an example image sequence of a buckling microtubule after washout. Buckling microtubules exhibit up to 12 µm long tubulin incorporation stretches that frequently extend along a large part of the microtubule lattice (**Fig. 3c,f; suppl.movies 5, 6**). In contrast, when keeping microtubules statically attached to motors using the non-hydrolysable ATP-analogue AMPPNP, we only observed tubulin incorporation stretches of less than 2 µm length (**Fig. 3d,f**). In gliding microtubules grown from biotin-free seeds, incorporation stretches also appear much shorter than in buckling microtubules (**suppl. movie 4**), although they are longer than in static microtubules (**Fig. 3e,f**), consistent with previous reports [9,11]. The spatial incorporation frequency also increases from AMPPNP to gliding and buckling microtubules, though the difference in incorporation frequency between buckling microtubules and the other two cases is relatively less pronounced (**Fig. 3g**). This may be due to overlapping incorporations that cannot be distinguished in our analysis (considering buckling microtubules exhibit very long incorporation stretches). Overall, we observed tubulin incorporation along 60% of the microtubule lattice length in buckling microtubules (**Fig. 3h**), five times as much as in gliding microtubules and 50 times as much as in static microtubules. Quantification of the total amount of tubulin turnover by accounting for both the longitudinal extent and the lateral spread of incorporation (**ED Fig. 6e**), we see that buckling microtubules show, by far, the highest turnover (**Fig. 3i**). While even unloaded motors were previously shown to induce microtubule damage and subsequent self-repair ^[9, 11, 33, 34]^, our observations reveal that kinesin-mediated buckling massively increases this effect.

**Figure 3:**
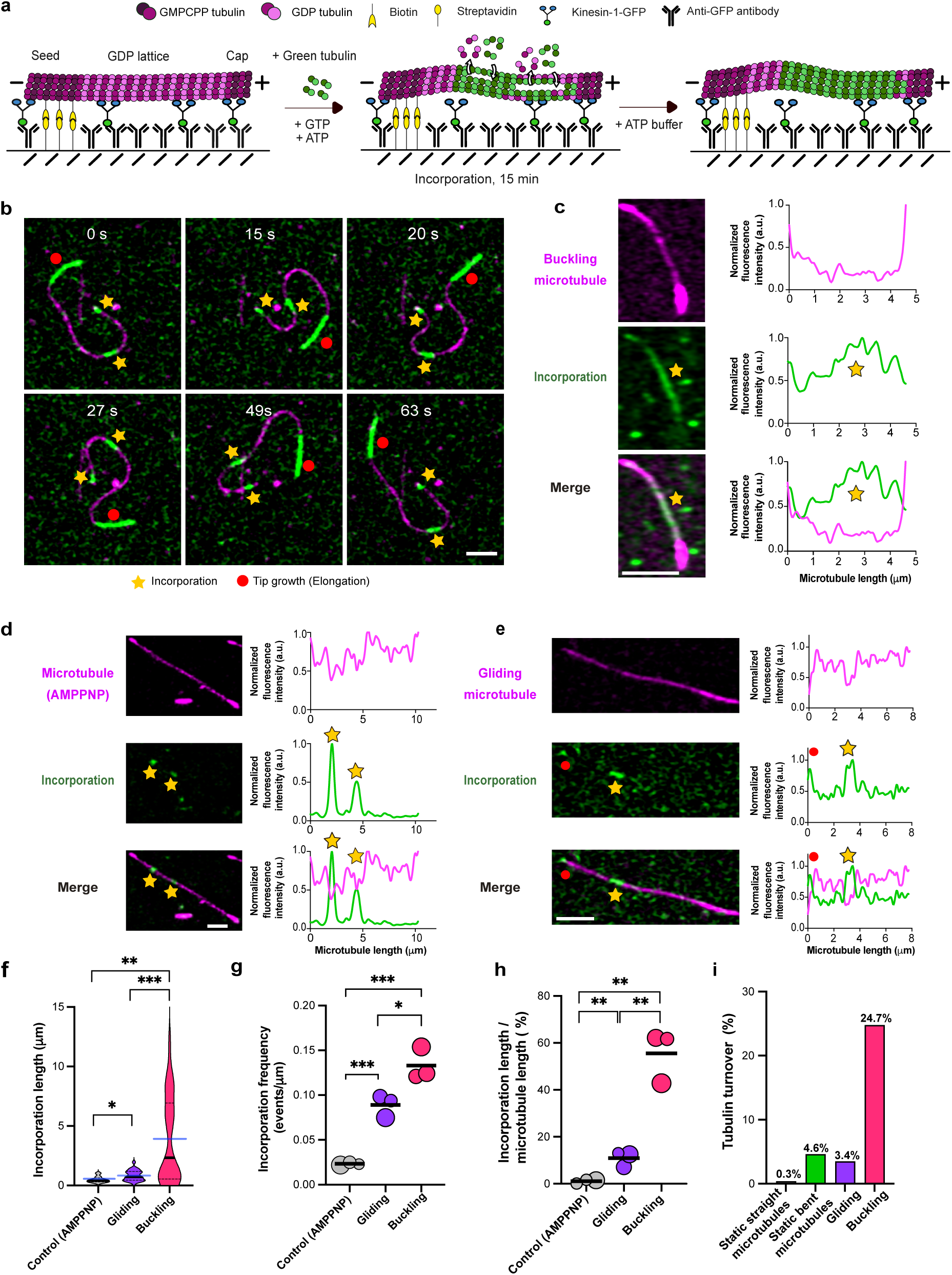
Extensive damage and consequent self-repair in buckling microtubules. **a**, Schematic of the experimental setup used to assess self-repair in buckling microtubules *in vitro*. **b**, Time-lapse sequence of a buckling microtubule showing incorporations of green labeled tubulin (marked with a yellow star) and the elongated tip (highlighted with a red circle). Scale bar: 2 μm. **c**, Example image showing incorporation of green labeled tubulin (marked with a yellow star) in a buckling microtubule. Scale bar: 2 μm. Graphs represent line scans of the microtubule (magenta) and the incorporation channel (green). Profiles have been normalized to 1 with the maximum value of the intensity (a.u.) of the microtubule and incorporation channel, respectively. Scale bar: 2 μm. **d**, Example image showing incorporation of green labeled tubulin (marked with a yellow star) in the buckling setup with static motors using AMPPNP. Graphs represent line scans of the microtubule (magenta) as well as the incorporation channel (green). Profiles have been normalized to 1 with the maximum value of the intensity (a.u.) of the microtubule and incorporation channel, respectively. Scale bar: 2 μm. **e**, Example image showing incorporation of green labeled tubulin (marked with a yellow star) in a gliding microtubule. Elongation of the microtubule tip is highlighted with a red circle. Graphs represent line scans of the microtubule (magenta) as well as the incorporation channel (green). Profiles have been normalized to 1 with the maximum value of the intensity (a.u.) of the microtubule and incorporation channel, respectively. Scale bar: 2 μm. **f**, Violin plots showing extensive incorporations in buckling microtubules. Total length of microtubules analyzed: 567 μm for gliding, 474 μm for buckling and 586 μm for control-AMPPNP from three independent experiments per condition. Black line represents the median and dotted lines represent the interquartile range. Blue lines represent the mean for each condition. p= 0.0018 (buckling-control), p= 0.0172 (gliding-control), and p= 0.0005 (gliding-buckling) using Mann-Whitney test (n= 9 incorporations for control-AMPPNP, n= 52 incorporations for gliding and n= 36 incorporations for buckling). **g**, Bubble plot showing a higher frequency of incorporations in buckling microtubules. Bubble sizes scale with the total microtubule length analyzed. Each circle represents an independent experiment (comprising of 226, 158 and 183 μm of total microtubule length analyzed for gliding; 142, 171 and 161 μm of total microtubule length analyzed for buckling and 218, 173 and 195 μm of total microtubule length analyzed for control-AMPPNP). Black line represents the mean. p= 0.0261 (for gliding-buckling); p= 0.0008 (gliding-control); p= 0.0005 (buckling-control) using unpaired-t-test. **h**, Bubble plot showing a higher amount of lattice length with incorporation, estimated as incorporation length/ microtubule length, in buckling microtubules *in vitro*. Bubble sizes scale with the total microtubule length analyzed. Each circle represents an independent experiment (comprising of 226, 158 and 183 μm of total microtubule length analyzed for gliding; 142, 171 and 161 μm of total microtubule length analyzed for buckling and 218, 173 and 195 μm of total microtubule length analyzed for control-AMPPNP). Black lines represent the mean. p= 0.0076 (gliding-control); p= 0.0010 (buckling-control) and p= 0.0026 (gliding-buckling) using unpaired t-test. **i,** Higher % of tubulin turnover in buckling microtubules. Total length of microtubules analyzed: 567 μm for gliding, 474 μm for buckling and 586 μm for control-AMPPNP from three independent experiments per condition. Refer methods for estimation of % tubulin turnover and **ED Fig. 6e** for estimation of amount of lateral tubulin incorporation.

### 4) Limits of microtubule self-repair under kinesin-induced buckling

The pronounced tubulin incorporation observed in buckling microtubules suggests that these microtubules experience substantial lattice damage. This leads to the question of whether microtubule self-repair is sufficient to counteract the damage sustained under such dynamic buckling. Since tubulin incorporation is the net outcome of damage and repair, we first assessed microtubule stability in the absence of free tubulin, where microtubules are known to spontaneously disassemble due to gradual tubulin loss even in the absence of external forces ^[8, 35]^. This condition serves as a reference point, removing the possibility of self-repair and allowing us to directly assess how strongly mechanical stress accelerates microtubule disassembly. We found that buckling microtubules frequently break and disassemble (**Fig. 4a**). Notably, we observed that breakage sites typically coincide with regions of high local curvature (**Fig. 4b; suppl video 7**), similar to what is seen in cells (**ED Fig. 3b** and in Odde et al., 1999 [27]). After just 4 min, 50% of the buckling microtubules had disassembled, and after 10 min, all buckling microtubules had disassembled (**Fig. 4c**). By contrast, gliding microtubules persist longer (as reported in Triclin et al., 2021 [9]): 50% remain intact after around 14 min, and within 30 min, all gliding microtubules had disassembled. When kinesin motors are rendered static using AMPPNP, microtubule survival is markedly enhanced, with 85 % of microtubules remaining intact even after 30 min.

**Figure 4:**
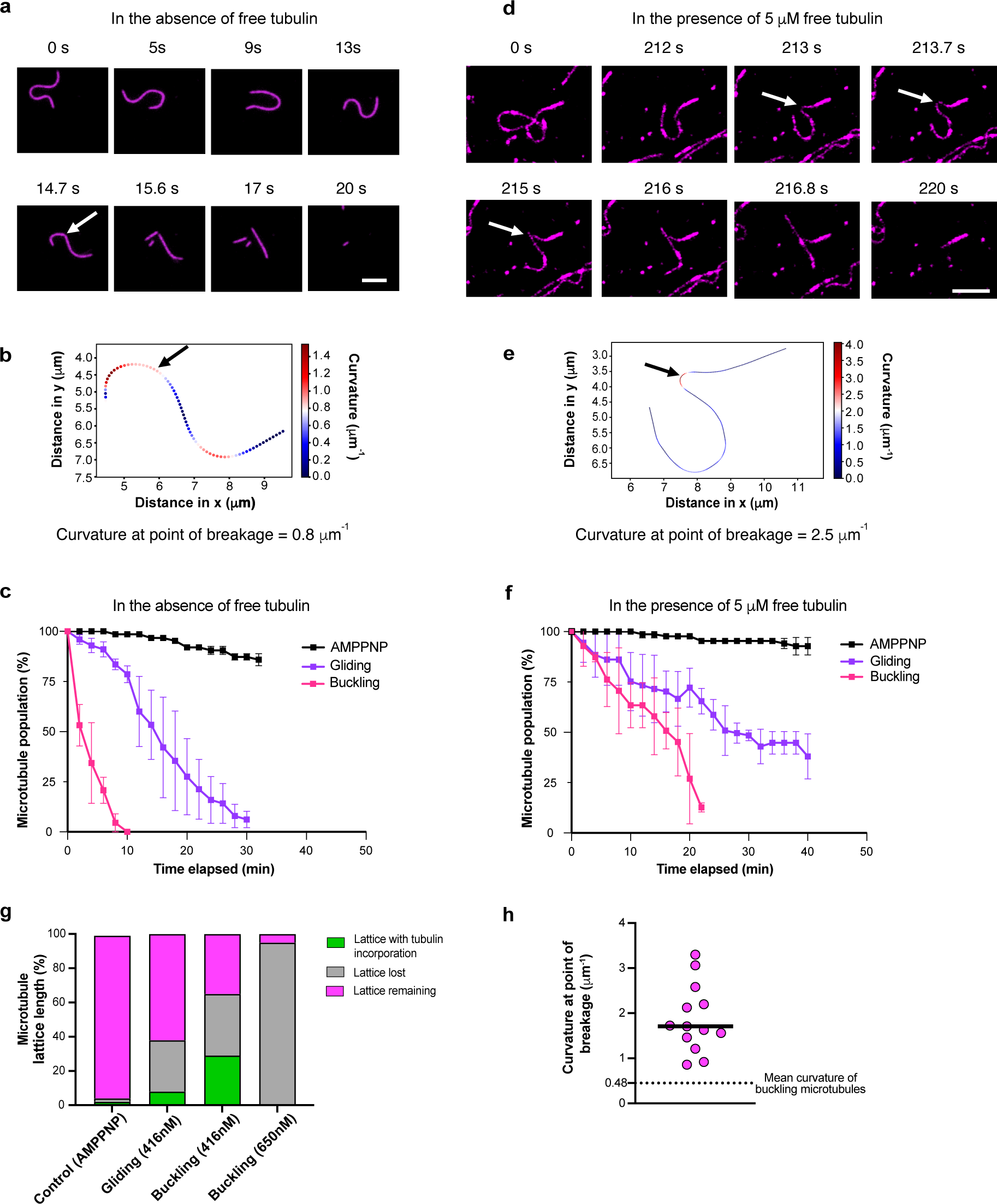
Motor-induced buckling damages microtubules beyond the limit of self-repair: **a**, Time-lapse sequence of a buckling microtubule breaking in the absence of free tubulin. White arrow in the frame at 14.7s indicate the point of breakage. Scale bar: 5 μm. **b**, Plot showing distribution of local curvature along the microtubule in Fig **4a** in the frame (at 14.7s) prior to microtubule breakage. Black arrow indicates point of breakage. **c**, Comparison of % microtubule population remaining in control (AMPPNP), gliding and buckling microtubules over time, in the absence of free tubulin. The symbols indicate mean ± S.D. Total length of microtubules analyzed: 567 μm for gliding, 474 μm for buckling and 586 μm for control-AMPPNP (no: of microtubules analyzed > 100) from two independent experiments per condition. **d**, Time-lapse sequence of a buckling microtubule breaking in the presence of 5 μM free tubulin. Point of breakage indicated by a white arrow. Scale bar: 5 μm. **e**, Plot showing distribution of local curvature along the trace of the microtubule in Fig. **4d** in the frame (at 213.7s) prior to microtubule breakage. Black arrow indicates the point of breakage. **f**, Comparison of % microtubule population remaining in control (AMPPNP), gliding and buckling assays over time in the presence of 5 μM free tubulin. The symbols indicate mean ± S.D. Total length of microtubules analyzed: 567 μm for gliding, 474 μm for buckling and 586 μm for control-AMPPNP (no: of microtubules analyzed >100) from two independent experiments per condition. **g**, Bar graph showing the relative proportion of microtubule lattice length lost, repaired (refer to Fig. 3 g,h) and remaining (refer Fig. 4f) at the end of 15 min, in the case of assays with AMPPNP, gliding microtubules (416 nM kinesin), buckling microtubules (416 nM kinesin), and buckling microtubules (650 nM kinesin). No: of microtubules analyzed > 100 from three independent experiments per condition. **h**, Curvature of buckling microtubules at point of breakage (n= 13 microtubules from eight independent experiments). Black line represents the median (1.71 μm^-1^). Black dotted line represents the average curvature of buckling microtubules (0.48 μm^-1^; Refer to Fig. 2g).

We next asked whether self-repair via tubulin incorporation could compensate for the damage observed under dynamic buckling (**Fig. 4d-f**). In the presence of free tubulin, gliding microtubules show markedly improved survival compared to the no-tubulin condition and statically-bound microtubules experience close to no loss (**Fig. 4f**), consistent with previous reports [9,11]. In contrast, buckling microtubules still disassemble rapidly, despite the availability of free tubulin (**suppl. movie 8**), typically after breakage in highly curved regions (**Fig. 4d,e,h; suppl. movie 7**). After just over 20 min, all buckling microtubules broke and disassembled (**Fig.4f**). Closer to physiological conditions, we used GDP microtubules in our assays. Our observations suggest that breakage is more frequent and occurs on shorter timescales than in studies with taxol-stabilized microtubules [36–38], which, by design were performed in the absence of free tubulin and did not account for self-repair.

To better understand the balance between damage and self-repair, we compared tubulin incorporation and microtubule survival after 15 min, in the presence of free tubulin across conditions (**Fig. 4g**). Microtubules with static motors (AMPPNP condition) show almost no detectable tubulin incorporation and remain nearly fully intact, indicating minimal lattice damage. In gliding microtubules, 30 % of the total initial microtubule length was lost due to breakage, and incorporation stretches appeared along 8 % of the initial lattice length. Buckling microtubules, driven by the same motor density used for gliding (416 nM), showed a greater loss of lattice length (49 %) and a much higher fraction of lattice length with incorporations (29 %). Strikingly, upon further increasing motor density (650 nM), microtubule lattice damage became catastrophic: 95 % of the initial length was lost, and no visible tubulin incorporation remained. This indicates that the rate of damage has surpassed the capacity for self-repair. Interestingly, the median value of curvature at point of breakage in buckling microtubules is 1.7 µm^-1^ (**Fig. 4h**), closely matching the value of 1.5 µm^-1^ reported by Odde et al.,1999 for occasional microtubule breakage in fibroblasts [27].

Together, these results highlight the limits of microtubule self-repair: Statically anchored microtubules remain stable, and gliding microtubules sustain moderate damage that is efficiently repaired, in line with previous reports that self-repair protects microtubules against breakage from motor motility [9]. In contrast, buckling – especially at high motor density – induces severe lattice disruption that exceeds self-repair capacity. Thus, dynamic buckling represents a mechanical challenge to microtubule integrity that cannot be balanced by intrinsic self-repair alone. Based on our experiments, breakage typically occurs in regions of high curvature, suggesting curvature as a determinant of lattice disruption. However, it remains unclear whether these curved regions are also subjected to elevated motor forces, since in our assays we cannot resolve local force patterns along the microtubule.

### 5) Worm-like chain model recapitulates microtubule buckling

To better understand the mechanical loads experienced by microtubules in our *in vitro* buckling assay, we developed a stochastic two-dimensional computational model that captures the key mechanical and dynamic features of our experimental system (**Fig. 5a**; see **SI simulation methods** for a detailed description). We modeled microtubules as worm-like chains composed of 8-nm segments – equivalent to the length of a tubulin dimer – the typical step-size of kinesin ^[39, 40]^. Microtubules are fixed at their minus ends, mimicking the anchored seeds in our experiments. Motors are distributed on a grid at variable densities and modeled as point-like entities that can bind to the nodes between microtubule segments and step along the microtubule (**Fig. 5a**). Since the motors remain attached to the substrate, their stepping leads to microtubule buckling. The motor parameters-including attachment and detachment rates and force-dependent stepping kinetics are based on literature values and our own measurements (see **ED Table 1**; **ED Fig. 4a-i**; SI simulation methods). Unless otherwise indicated, we performed simulations using 10 µm-long microtubules with 5 mm persistence length (L_p_) and a motor density (ρ) of 400 µm^-2^. Note that the model does not rely on any fit parameters tuned to our experimental system. Rather than aiming for a precise quantitative match, we chose a coarse-grained model designed to semi-quantitatively capture essential trends. To verify that the model reproduces our experimental observations, we compared the mean microtubule curvature between simulation and experiments (**ED Fig. 5a**).

**Figure 5:**
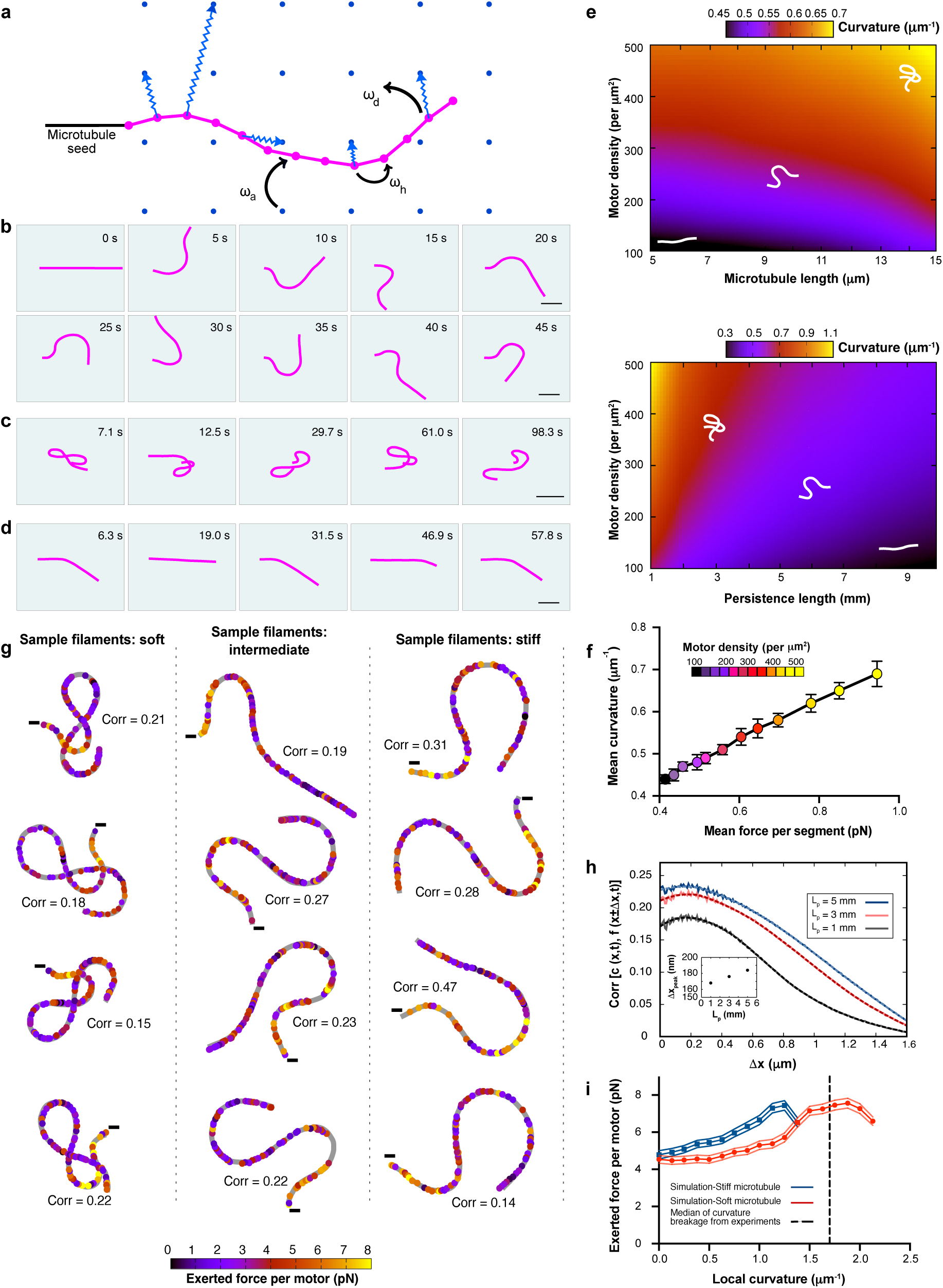
Worm-like chain model-based simulations capture microtubule deformation under active forces: **a**, Schematic of the simulation setup: motor proteins (represented as blue dots) are uniformly distributed on the substrate and interact with the microtubule by attaching, hopping along it, or detaching with rates ω_a_, ω_h_ and ω_d_ respectively. The minus end of the microtubule (seed, represented in black) is kept fixed. Springs represent pulling forces exerted by attached motors. **b-d**, Representative microtubule conformations in different deformation regimes: **b**, regular flagella-like oscillations; **c**, loop formation; and **d**, regular beating. The motor density is ρ= 248, 494 and 123 μm^−2^, respectively. Scale bars: 2 μm. Other parameters are set to default values mentioned in ED Table 1. **e**, Numerical phase diagrams of microtubule deformation under active motor forces. Mean curvature of microtubules as a function of (top) microtubule length and motor density at a fixed persistence length L_p_= 5 mm and (bottom) persistence length and motor density at a fixed microtubule length L= 10 μm. The heat maps display ensemble-averaged data over 10^5^ filaments per parameter set. The underlying data are discretized into 20 bins along each axis, and a Gaussian smoothing filter was applied to generate the final color map. Schematic white filaments show representative microtubule configurations for corresponding mean curvature values. **f**, Mean curvature of the filament versus the mean force exerted on the filament per segment for L = 10 µm, L_p_ = 5 mm, and increasing motor density. The data is ensemble averaged over 10^3^ filaments. **g**, Representative microtubule configurations from simulations, color-coded by the local active force exerted by molecular motors. Each dot represents an attached motor. Shown are examples for three levels of bending rigidity: soft filaments with L_p_ =1 mm (left), intermediate stiffness with L_p_ = 5 mm (middle), and stiff filaments with L_p_ = 10 mm (right). Other parameters: L = 10 μm, ρ = 494 μm^−2^. In each panel, the gray line traces the filament backbone. Colored circles mark the positions of the active motors along the microtubule, with the color indicating the magnitude of the exerted force by each motor. Minus signs denote the fixed ends of the microtubules. 10−15% of the modeled microtubules segments are occupied by motors (refer to **ED Fig. 5b**). Configurations highlight differences in deformation patterns and force localization depending on filament stiffness. For each case, the corresponding curvature-force correlation is also shown, which is computed at the optimal spatial offset (within the narrow range [0.17, 0.20] μm) and averaged along the filament. **h**, Cross-correlation between the local curvature c(x,t) of a modeled buckling microtubule at position x and time t, and the local force f(x±Δx,t) exerted at spatially offset positions at the same time, plotted as a function of spatial offset Δx (See SI simulation methods for details of Pearson correlation calculation). Results are shown for varying persistence lengths from an ensemble of 10^5^ microtubules. Solid lines represent raw simulation data, while dashed lines indicate smoothed curves obtained using a Savitzky-Golay filter. Default values for all other parameters are listed in ED Table 1. Cross-correlation values are averaged over time and along the length of the filament. Inset shows the dependence of the correlation peak position on persistence length (L_p_). **i,** Force exerted by each motor versus local curvature, shown for simulated stiff microtubule with L_p_ = 10 mm (dark blue) and soft microtubule with L_p_ = 1 mm (red). Other parameters: L = 5 μm, ρ = 400 μm^−2^. Solid lines represent the mean, and shaded regions indicate standard deviations across the ensemble of 10^5^ microtubules. Vertical dotted line (in black) represents the median curvature of breaking buckling microtubules (1.7 µm^-1^) from Fig. 4h.

By varying the motor density, we were able to reproduce the distinct dynamic behaviors observed in our experiments: flagella-like oscillations (**Fig. 5b**, compare to Fig. 2d; **suppl. movie 9**), looping (**Fig. 5c**, compare to Fig. 2e, **suppl. movie 10**), and beating motions (**Fig. 5d**, compare to Fig. 2f; **suppl. movie 11**). We then systematically explored the dependence of microtubule curvature on key parameters by constructing phase diagrams showing the mean microtubule curvature as a function of motor density, microtubule length, and microtubule persistence length (**Fig. 5e**). This reveals intuitive trends: curvature increases with higher motor density and microtubule length and decreases with increasing stiffness.

To assess how motor-generated forces influence microtubule shape, we then examined how mean curvature depends on the mean force along the microtubule. **Fig. 5f** shows the mean microtubule curvature as a function of the mean force per microtubule segment at varying motor densities, which effectively increases the number of active motors on the microtubules (**ED Fig. 5b**). Mean curvature scales predictably with the overall applied force.

Next, we aimed to determine which local motor force patterns are responsible for inducing buckling. **Fig. 5g** shows color-coded force profiles for twelve representative microtubules with different persistence lengths (L_p_ = 1, 5 and 10 mm). Notably, the force distribution is highly heterogeneous and does not consistently coincide with regions of high curvature. When correlating local curvature with the local force per segment, we found only a modest correlation (r ≈ 0.2). This likely reflects the inherent complexity of the system, in which distributed and competing active forces act over and are transmitted along a filament with internal degrees of freedom.

To further investigate the relationship between force application and resulting curvature, we calculated the correlation between local force and curvature as a function of the spatial offset Δx between the force application point and the measured curvature. The correlation between local force and curvature is generally weak, with a peak at Δx ≈ 0.2 µm (**Fig. 5h**), suggesting that curvature tends to arise slightly displaced with respect to the site of force application. This behavior can be intuitively understood with an analogy to a flexible rod that is exposed to compression: the resulting bend is not maximal at the point of force application but depends on how the rod distributes stress along its length. Soft microtubules show weaker correlations that decay more strongly with distance Δx compared to stiff microtubules (**Fig. 5h**). This is consistent with their higher susceptibility to noise, which makes their mechanical response less deterministic – also evident in their chaotic looping behavior (**Fig. 5e, bottom**).

Finally, we examined how local motor forces relate to microtubule breakage. In our experiments, microtubules consistently break at regions of high curvature (Fig. 4h), similar to what is seen in cells (Fig. 2b, bottom; [24]). Our measurements show that curvature at sites of breakage is three times as high as the mean curvature of buckling microtubules (1.7 µm^-1^ compared to 0.48 µm^-1^, see Fig. 4h). Consistent with the low correlation between force and curvature, our simulations reveal that for a given curvature, the force per microtubule segment varies substantially (see **ED Fig. 5c** for a soft microtubule and **ED Fig. 5d** for a stiff microtubule). Nevertheless, a clear trend emerges: higher local curvatures tend to be related to a slightly higher mean force per motor, both in soft and stiff microtubules (**Fig. 5i**). Notably, the model predicts the largest forces to be associated with curvatures in the range of 1 µm^-1^ (for stiff microtubules) to 2 µm^-1^ (for soft microtubules), which coincides with the curvature range where microtubule breakage is observed experimentally (median curvature at breakage is indicated by the dashed line in **Fig. 5i**, also refer Fig. 4h).

Interestingly, the force exerted on each segment of the filament displays a nonmonotonic relationship with the local curvature (**Fig. 5i**). At low curvatures, the curvature increases gradually with force. As deformation progresses, the system enters a nonlinear force-curvature regime characterized by an accelerated increase in curvature [41]. At very high curvatures, however, the force begins to decline, coinciding with topological transitions in the filament, such as the formation of loops. In this regime, the force required for further deformation may decrease, driven by abrupt, large-scale conformational changes of the filament. Overall, the model reveals that local curvature and local motor-generated forces only weakly correlate and are not linked in a simple manner.

### 6) Microtubule damage arises from a combination of curvature and motor action

To summarize our findings so far, we have established that: (i) Static curvature is sufficient to induce microtubule damage and self-repair, but it neither leads to microtubule breakage nor reaches the levels of tubulin turnover observed in cells. (ii) Dynamic buckling, in contrast, causes extensive lattice disruption and frequent breakage, which typically occurs in regions of high curvature. (iii) Modeling reveals that local curvature and local motor-induced force patterns correlate only moderately, yet overall higher forces give rise to higher mean curvatures. Together, these observations indicate that curvature alone cannot account for the severe disruption seen during buckling, and suggest that curvature acts in concert with motor motility [9] and motor-induced forces to damage microtubules. To experimentally disentangle the contribution of force from curvature, we modified our buckling assay to isolate the effects of force. Specifically, we included biotinylated tubulin in both microtubule ends (seed and cap), allowing microtubules to be anchored at both ends (**Fig. 6a**). This prevents buckling and keeps microtubules straight but still allows surface-attached motors to exert pulling forces on the microtubule lattice. We will refer to these microtubules as “double-anchored microtubules” hereafter.

**Figure 6:**
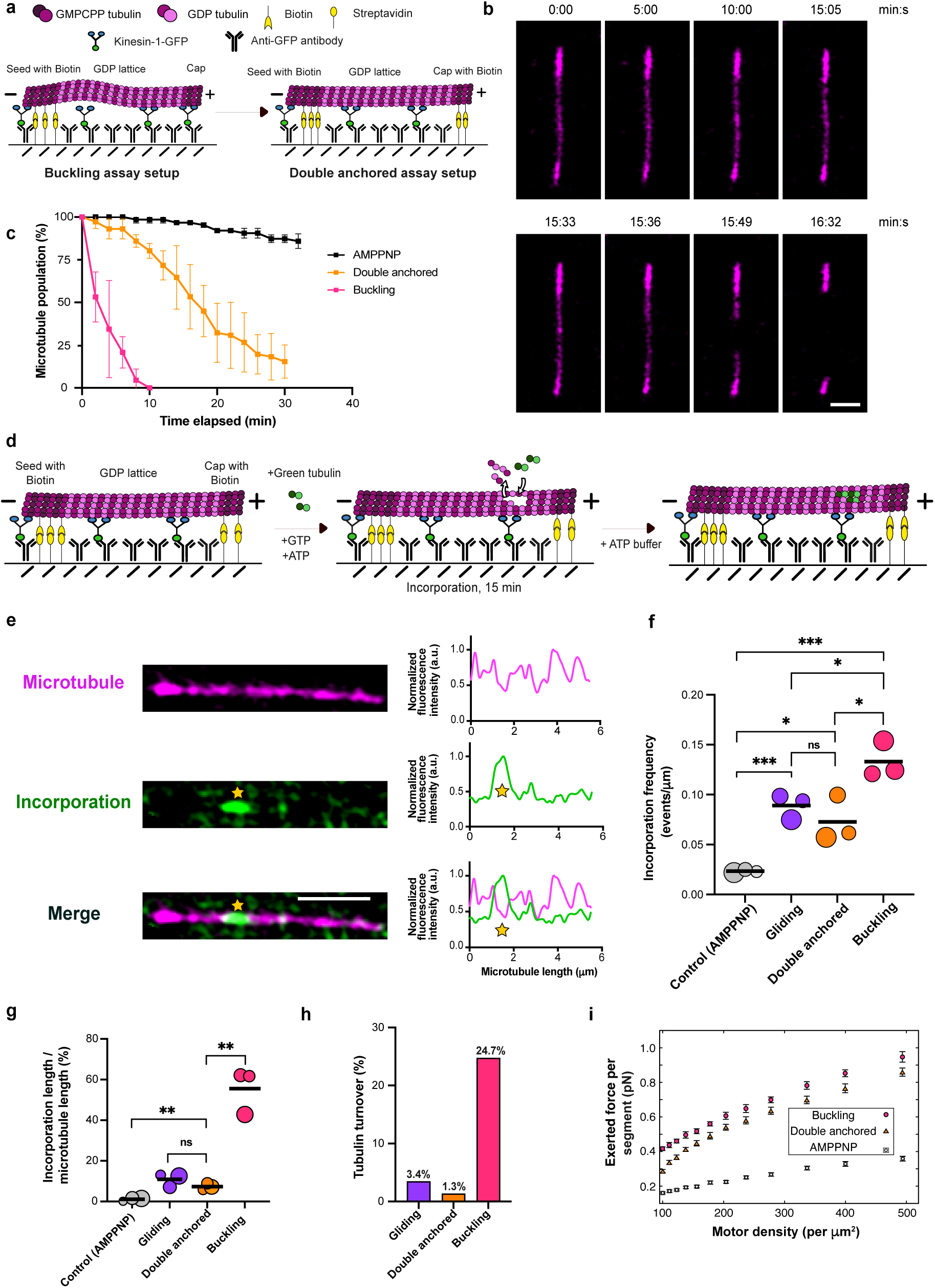
Combination of curvature and motor action contribute to extensive damage in buckling microtubules: **a**, Schematic of the experimental setup used to subject microtubules to pulling forces of kinesin (double-anchored assay). In this assay, both ends of the capped microtubule are anchored to the surface via biotin/streptavidin. **b**, Time-lapse sequence of a double-anchored microtubule subjected to motor-induced pulling forces in the absence of free tubulin. Scale bar: 2 μm. **c**, Comparison of % microtubule population remaining in control (AMPPNP), double anchored and buckling assays over time in the absence of free tubulin. The symbols indicate mean ± S.D. (n> 100 microtubules analyzed in each condition from two independent experiments) **d**, Schematic of the experimental setup used to assess self-repair in double-anchored microtubules. **e,** Example image showing incorporation of green labeled tubulin (marked with a yellow star) in a microtubule (magenta) in the double-anchored assay. Scale bar: 2 μm. Graphs represent line scans of the microtubule (magenta) as well as the incorporation channel (green). Profiles have been normalized to 1 for the maximum value of the microtubule and incorporation channel, respectively. **f**, Bubble plot showing lower frequency of incorporations in double-anchored microtubules compared to buckling microtubules. Bubble sizes scale with the total microtubule length analyzed. Each circle represents an independent experiment. Data from three independent experiments comprising 98, 139 and 110 μm of the total microtubule length analyzed for double anchored microtubules. Refer Fig. 3g for details on total microtubule length analyzed for gliding and buckling microtubules. Black lines represent the mean. p= 0.0242 (for double-anchored-buckling); p= 0.0212 (double-anchored -control); p= 0.3432; not significant (double-anchored-gliding) using unpaired t-test. **g**, Bubble plot showing a lower percentage of lattice length with incorporation, estimated as incorporation length/ microtubule length, in the double-anchored assay when compared to buckling as well as control (AMPPNP). Bubble sizes scale with the total microtubule length analyzed. Each circle represents an independent experiment (comprising 98, 139 and 110 μm of total microtubule length analyzed for double anchored microtubules; Refer Fig. 3h for details on total microtubule length analyzed for gliding and buckling microtubules). Black lines represent the mean. p= 0.0017 (double anchored -buckling) and p= 0.1415, not significant (double anchored -gliding); p= 0.0077 (double anchored -control-AMPPNP) using unpaired t-test. Refer Fig. 3h for the p-values comparing control (AMPPNP), gliding and buckling. **h**, Lesser % of tubulin turnover in double anchored microtubules when compared to buckling microtubules. Total length of microtubules analyzed: 567 μm for gliding, 347 µm for double anchored and 474 μm for buckling microtubules from three independent experiments per condition. Refer to the methods for estimation of % tubulin turnover and ED Fig. 6e for estimation of amount of lateral tubulin incorporation. **i**, Effect of motor protein arrangement and mobility for three configurations: (1) motors distributed across the surface (leading to buckling), (2) motors arranged linearly beneath an straight microtubule (mimicking double-anchored microtubules), and (3) in presence of AMPPNP. Total forces exerted on each discretized node of the microtubule are plotted as functions of motor density. Parameters: L = 10 μm and L_p_ = 5 mm. Data are averaged over time and across five microtubules.

**Fig. 6b** shows an example of such a double-anchored microtubule in the absence of free tubulin. Despite remaining straight, the microtubule eventually breaks and disassembles after approx. 16 min (**suppl. movie 12**). A survival analysis (**Fig. 6c**) confirms that double-anchored microtubules are significantly more stable than buckling ones, though much less so than static controls without active motors.

We reasoned that the reduced damage observed in the absence of free tubulin should result in less tubulin incorporation in the presence of free tubulin. To test this, we added free, green-labeled tubulin to the assay for 15 min (**Fig. 6d**). Double-anchored microtubules show occasional incorporation of free tubulin, as seen in the example in **Fig. 6e**. Quantification (**Fig. 6f-h; ED Fig. 6a,e**) reveals that incorporation levels are slightly elevated compared to static microtubules, but remain significantly lower than those observed in buckling microtubules. This indicates that double-anchored microtubules undergo limited damage and self-repair, consistent with their increased stability in the absence of free tubulin, when compared to buckling microtubules.

To determine whether these differences could be explained by differing motor forces, we compared the mean motor-generated force per microtubule segment between double-anchored and buckling microtubules in simulations across a range of motor densities (**Fig. 6i; ED Fig. 6b,c**). The results show that double-anchored microtubules experience forces of comparable magnitude to buckling microtubules – particularly at intermediate and high motor densities, as used in our experiments – and far greater than in static bent (**ED Fig. 6d**) as well as AMPPNP conditions.

In conclusion, despite being subjected to similar levels of motor-generated forces, only buckling microtubules exhibit extensive damage and self-repair. The comparable levels of tubulin loss and incorporation in gliding and double-anchored microtubules further suggest that motor motility, in addition to motor-generated forces, contributes to lattice disruption. Thus, lattice damage likely arises from motor movement (and the forces associated with it) and is markedly amplified in zones of high curvature. In summary, our observations support the view that damage in buckling microtubules stems from a combination of curvature and motor action, with force playing a surprisingly minor role and curvature acting as the key amplifier of motor-induced damage.

### 7) Intracellular factors enhance microtubule resilience to buckling-induced damage

The extensive damage observed in buckling microtubules, which often surpasses their intrinsic self-repair capacity, prompted us to explore how microtubules withstand mechanical stress in the intracellular environment – where bending and buckling are common, as in our *in vitro* system. Several microtubule-associated proteins (MAPs) have been implicated in promoting microtubule resilience to mechanical stress[13,42–44]. We therefore hypothesized that intracellular factors may protect microtubules from damage and help maintain their structural integrity.

To test this idea, we examined the survival of buckling microtubules in the presence of cell lysate from HEK293 cells, but in the absence of free tubulin (**Fig. 7a**). While the exact composition of the lysate is unknown, it contains soluble cytoplasmic components and could thus influence microtubule stability. Because cell lysate has previously been shown to modulate motor activity [45], we used a low concentration of 20 µg ml^-1^ to minimize such effects, as shown in Korten et al., 2013 [46]. Using a low concentration of cell lysate also ensures that any residual tubulin present is negligible. We tested the motor activity of purified kinesin both in the absence and in the presence of 20 µg ml^-1^ of HEK293 lysate and found the microtubule gliding velocity to be similar **ED Fig. 7a**). An example of a buckling microtubule in the presence of lysate is shown in **Fig. 7b**.

**Figure 7:**
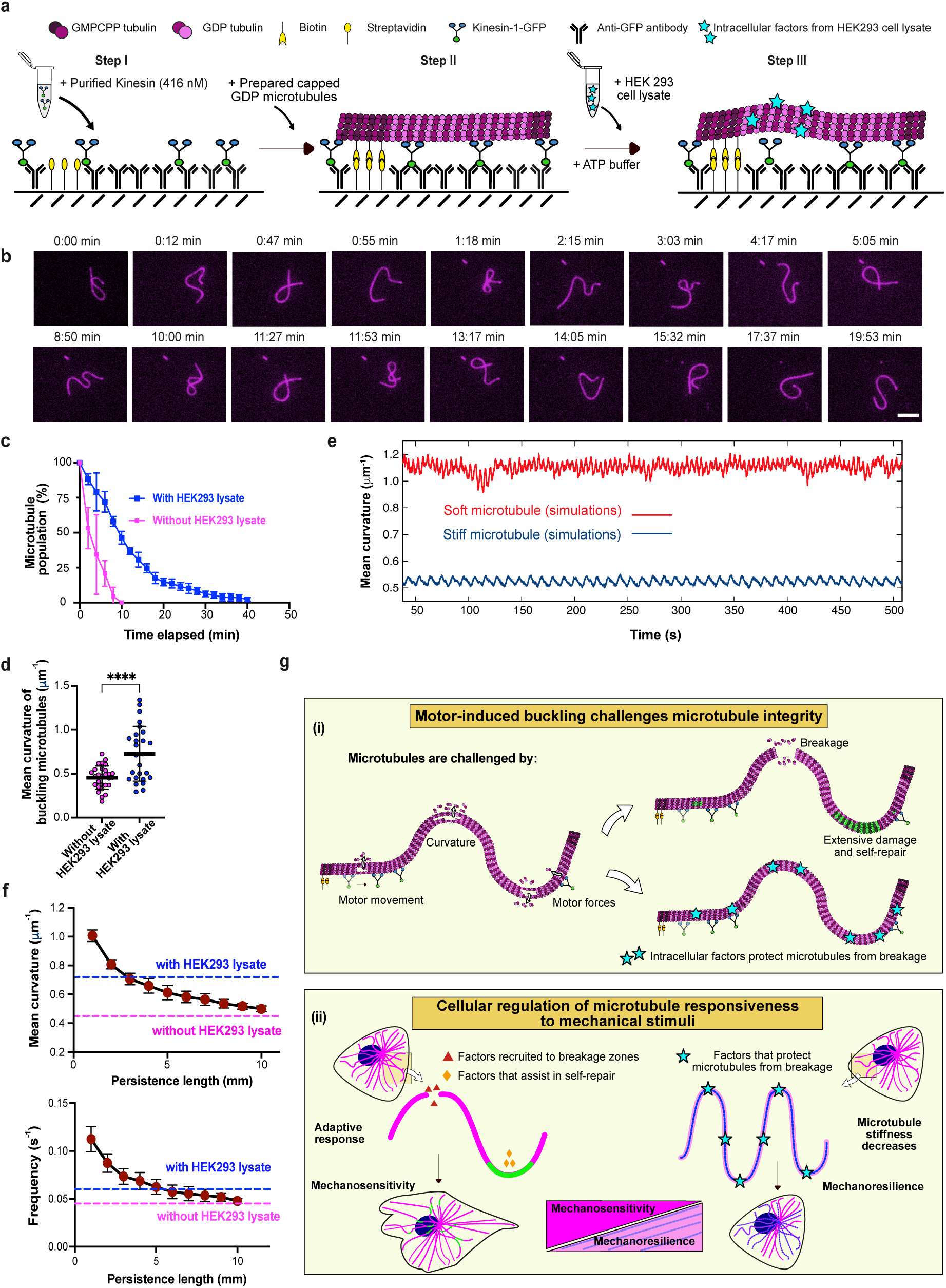
Intracellular factors protect buckling microtubules from breakage: **a**, Schematic of the experimental setup used to assess the influence of intracellular factors on microtubule survival in buckling assays: In step I, similar to the gliding and buckling assay setups, purified kinesin-1-GFP (416 nM) is immobilized on to an anti-GFP antibody coated surface. Then, prepared capped microtubules (step II) are added. Microtubule buckling (step III) is achieved by flushing in a mix containing ATP buffer (GTP, ATP) with 20 µg ml^-1^ HEK293 cell lysate. **b**, Time-lapse sequence showing survival of a buckling microtubule in presence of 20 µg ml^-1^ HEK293 cell lysate. Scale bar: 5 μm. **c**, Buckling microtubules survive longer in presence of cell lysate. Comparison of the percentage of the microtubule population remaining (in the absence of free tubulin) over time in the case of buckling microtubules both with and without 20 µg ml^-1^ HEK293 lysate (n> 100 microtubules analyzed in each condition from two independent experiments). The symbols and error bars indicate mean ± S.D respectively. **d**, Comparison of mean curvature in buckling microtubules in the presence and absence of HEK293 cell lysate. Black line represents the mean and error bars represent the S.D. p<0.0001 using an unpaired t-test. (Mean obtained from analyzing microtubule mean curvature at different timepoints from 4 microtubules in each case: n= 28 for without HEK293 lysate condition and n= 26 for with HEK293 lysate condition. Refer **ED Fig. 7b** for comparison of lengths of microtubules analyzed). **e**, Effect of microtubule stiffness on mean curvature. Examples of mean curvature evolution over time in simulations. The behavior of a soft microtubule (in red) with L_p_ = 1 mm is compared to a stiff microtubule (in dark blue) with L_p_ = 10 mm (slight bending phase). Other parameter values are L = 10 μm and ρ = 400 μm^−2^. **f**, Mean curvature (top) and oscillation frequency (bottom) as functions of microtubule persistence length at L = 10 μm and ρ = 400 μm^−2^. The symbols and error bars indicate mean ± S.D respectively. Blue and pink dotted lines represent experimentally determined values with and without HEK293 lysate respectively. **g (i)** Buckling microtubules are challenged by the combination of motor motility, forces and curvature resulting in extensive damage and self-repair or breakage. Intracellular factors protect microtubules and help enhance their survival under sustained deformation. **g(ii)** Our findings suggest that cells may use intracellular factors to regulate microtubule response to mechanical stimuli by factors specifically recruited to microtubule breakage zones and repair sites (mechanosensitivity) and by factors that decrease microtubule stiffness (mechanoresilience).

We observed that the addition of lysate during buckling increases microtubule survival: buckling microtubules remain intact for up to ∼40 min, compared to ∼20 min without lysate (**Fig. 7c**). Since no free tubulin was added, this enhanced survival suggests that factors present in the lysate reduce tubulin loss and stabilize the microtubule lattice. Interestingly, when comparing buckling microtubules in the absence and presence of lysate (**Fig. 2d vs. 7b**; suppl. movie 13), we noticed that microtubules visually appear softer in the lysate condition, since they buckle faster and with more pronounced curvature. This impression is supported by quantification: buckling microtubules exhibit higher mean curvatures in the presence of lysate (**Fig. 7d**).

To better understand these observations, we used our computational model to compare the buckling behavior of soft (L_p_ = 1 mm) and stiff (L_p_ = 10 mm) microtubules. Soft microtubules show both a higher buckling frequency and greater mean curvature (**Fig. 7e**), resembling the experimental data in the presence of lysate. The experimental difference can be captured by an approximate two-fold reduction in the persistence length in the presence of lysate (**Fig. 7f**). Alternatively, intracellular factors could modulate motor motility parameters. Model predictions indicate that reproducing the lysate data would require either a 2-7-fold increase in the attachment rate ω_a_ (**ED Fig. 7c**) or a roughly two-fold decrease in the detachment rate ω_d_ (**ED Fig. 7d**). Importantly, both scenarios increase the effective number of motors attached and thus cannot explain the higher survival in lysate, which would require the opposite effect (fewer attached motors). We therefore consider changes in motor parameters alone unlikely to account for the experimental data, unless additional protective factors overcompensate the increased damage.

Although we cannot currently identify the molecular players responsible for this effect, our findings support the interpretation that cells may promote microtubule survival under mechanical stress primarily by enhancing microtubule flexibility. Increased flexibility may allow microtubules to adopt strongly bent conformations without breaking, thus preserving lattice integrity under sustained deformation.

## Discussion

In this study, we highlight the limits of intrinsic microtubule self-repair under mechanical stress induced by molecular motors and show that additional intracellular factors are required to maintain microtubule integrity during sustained dynamic buckling. Using an *in vitro* assay that mimics kinesin-driven buckling as observed in cells, we find that the combination of motor motility, motor-induced forces and high curvature inflicts substantial lattice damage that often exceeds microtubule self-repair capacity. While microtubules intrinsically respond to this damage by incorporating free tubulin, these repair events are often not sufficient to prevent breakage. In contrast, the presence of intracellular components markedly enhances microtubule resilience, enabling microtubules to survive under mechanically demanding conditions (**Fig. 7g(i)**).

Among the cytoskeletal filament types, microtubules stand out due to their unusual combination of properties: they are relatively stiff polymers, yet remarkably responsive to biochemical and mechanical stimuli [3,14,18]. Since the discovery of microtubule self-repair, it has been widely assumed that this mechanism alone is sufficient to counteract the mechanical forces and damage they sustain in cells [6,9,14,15]. However, our results challenge this notion. They reveal that in situations where microtubules are subjected to dynamic buckling by continuous motor activity, leading to high curvatures, as seen in cells, intrinsic self-repair mechanisms can be overwhelmed.

Interestingly, microtubules show higher curvatures in the presence of cell lysate than in buffer alone, suggesting that intracellular factors may stabilize microtubules by modulating their mechanical properties. While changes in motor parameters such as altered attachment or detachment rates could also contribute to the observed buckling behavior, such changes alone would not readily account for the increased survival of microtubules in the presence of cell lysate. This observation therefore raises the possibility that microtubule stiffness in cells may be lower than previously estimated from *in vitro* measurements [18,47,48]. Reduced stiffness could serve as a protective function by allowing microtubules to adjust their shape under load – a form of mechanical compliance that may potentially be actively regulated in cells. For example, cells may selectively increase microtubule flexibility through recruitment of MAPs that reduce microtubule rigidity [49,50]. Tubulin post-translational modifications, particularly acetylation, have been proposed to enhance microtubule flexibility by weakening the lateral interactions between tubulin dimers [51]. However, in our experiments, it is unlikely that acetylating enzymes from the cell lysate contribute significantly to the observed microtubule resilience, given the short timescale [52].

More broadly, our findings suggest the intriguing possibility that cells may fine-tune microtubule rigidity and responsiveness to mechanical signals depending on functional needs (**Fig. 7g(ii)**). For example, primary cilia, which serve as sensory organelles, may benefit from the ability of microtubules to respond to subtle mechanical stimuli (mechanosensitivity), whereas motile cilia may require less sensitive microtubules to withstand repetitive mechanical stress (mechanoresilience). In plant cells, highly organized microtubule arrays guide morphogenesis by aligning with tensile stress patterns, often forcing them into strongly curved conformations [53]. By differentially regulating stiffness and stress responsiveness, cells may thus navigate the conflict between cell geometry and tension patterns. A better understanding of how microtubule properties are regulated in different contexts could reveal cell-type specific adaptations and vulnerabilities.

## Materials and Methods

### Cell culture and live cell imaging

Male potoroo kidney epithelial cells (PtK2), stably expressing GFP-Tubulin (displayed in figure 2 in Magenta) were cultured at 37°C and 5% CO_2_ in DMEM/F12 media (31331028, Gibco), supplemented with 10% Fetal Bovine Serum (FBS, Gibco A5670701) and 1% Penicillin-Streptomycin solution (15070063, Gibco). Human Embryonic Kidney cells (HEK293, DMSZ, ACC305) were cultured in Complete DMEM (high glucose with HEPES) media (Fisher Scientific, 42430025) with 10% Fetal Bovine Serum (heat inactivated at 56°C) and 1% Penicillin-Streptomycin solution at 37°C and 5% CO_2_. For live cell imaging, PtK2 cells were seeded in confocal glass-bottom dishes (734-2904, VWR Avantor) coated with 0.01 mg ml^-1^ fibronectin, a day prior to imaging. Live cell imaging for observing dynamic microtubules was carried out using a 63x immersion oil objective (Zeiss) maintained at 37°C and 5% CO_2_. Time-lapses of bending and buckling events in PtK2 cells were captured for a period of 5 min.

### Purification and labelling of tubulin

Tubulin free of Microtubule Associated Proteins (MAPs) was purified from fresh calf brains by three cycles of polymerization-depolymerization, using a combination of low and high salt buffers, followed by cation exchange chromatography, as previously described in [54]. Briefly, the first polymerization-depolymerization cycle was performed in low salt conditions (0.1 M PIPES, 0.5 mM MgCl_2,_ 2 mM EGTA and 0.1 mM EDTA). This was followed by the second cycle in high salt buffer [High Molarity PIPES Buffer: 1 M PIPES, pH 6.9, supplemented with KOH, 10 mM MgCl_2_, 20 mM EGTA]. Subsequently MAP free tubulin was obtained after cation-exchange chromatography (Fractogel EMD SO3, Merck) in 50 mM PIPES, pH 6.8, supplemented with 1 mM MgCl_2_ and 1 mM EGTA.

Fluorescently labeled tubulin (ATTO-488 and ATTO-565-labeled tubulin) and biotinylated tubulin were prepared according to the protocol described in Hyman et al.,1991 [54]. MAP-free tubulin obtained after cation exchange chromatography was polymerized at 37°C for 1 hour in BRB80 [Brinkley Buffer 80: 80 mM PIPES, pH 6.8, 1 mM EGTA and 1 mM MgCl_2_] supplemented with 33% glycerol, 4 mM MgCl_2_ and 1 mM GTP. The polymerized microtubules were layered onto prewarmed cushions of 0.1 M Na-HEPES, pH 6.8, 1 mM MgCl_2_, 1 mM EGTA, 60% v/v glycerol. This was followed by high-speed centrifugation at 37°C for 1 hour. Then the pellet was resuspended in 0.1 M Na-HEPES, pH 8.6, 1 mM MgCl_2_, 1 mM EGTA, 40% v/v glycerol. To this, 1/10^th^ volume of 100 mM NHS-ATTO-565/488 (ATTO-Tec), or NHS-LC-LC-biotin (EZ-link, ThermoFisher) was added, and the mix was allowed to incubate for 10 min at 37°C. Subsequently, the labeling reaction was stopped using two volumes of BRB80, with 100 mM potassium glutamate and 40% v/v glycerol. The labeled microtubules were sedimented onto BRB cushions with 60% glycerol. Following this, an additional cycle of polymerization and depolymerization was performed before the labeled tubulin was aliquoted, snap-frozen and stored in liquid nitrogen till use.

### Purification of Kinesin-1-GFP motor proteins

Recombinant, truncated kinesin-1 motor protein was purified as previously described in [55]. The plasmid encoding for the kinesin-1 protein (truncated to 560 aa), pET17_K560_GFP_His, was purchased from Addgene (15219, Cambridge, MA). The plasmid was transfected into Rosetta2 (DE3)-pLysS *E. coli* (VWR) and incubated with 0.2 mM IPTG at 16°C for 16 h. Following this, the cells were resuspended in cation-exchange buffer (6.67 mM Na-Acetate, 6.67 mM MES, 6.67 mM HEPES, 20 mM β-mercaptoethanol (BME), 0.2 mM ATP, and 0.2% Tween-20, pH 7.0 supplemented with a protease inhibitor cocktail) and lysed by sonication. The lysates were centrifuged at 38,000xg for 30 min at 4°C and loaded into a HiTrap SP Cation exchange column (HiTrap SP HP™, 17-1151-01, Cytiva). The column was then washed with cation exchange buffer supplemented with 50 mM KCl, followed by elution with cation exchange buffer supplemented with 300 mM KCl. The eluted fraction was then loaded on to a Ni-NTA column (HisTrap HPTM, 17-5247-01, Cytiva) after dilution with nickel loading buffer (Nickel buffer: 50 mM sodium phosphate buffer, pH 7.5, 5% w/v glycerol, 300 mM KCl, 1 mM MgCl_2_, 0.2% w/v Tween-20, 10 mM BME, 0.1 mM ATP, supplemented with imidazole to a final concentration of 36 mM). This was followed by washing with nickel washing buffer (nickel buffer supplemented with KCl to a final concentration of 1000 mM and imidazole to a final concentration of 30 mM) and elution with nickel elution buffer (nickel buffer supplemented with imidazole to a final concentration of 300 mM). The collected fractions were centrifuged at 4,000 g for 30 minutes at 4 °C. To remove imidazole, the sample was dialyzed overnight against K560 buffer (50 mM Na-phosphate pH 7.5, 300 mM KCl, 5% glycerol, 1 mM MgCl₂, 1 mM DTT, 0.1 mM ATP). The sample was then loaded onto a gel filtration Superdex column (Superdex 200 Increase 10/300 GL, 28-9909-44, Cytiva) and eluted with K560 buffer. The eluted fractions were concentrated using a 30 kDa membrane filter by centrifugation at 4,000xg for 30 minutes at 4°C. Finally, the protein was snap-frozen in 5 µl aliquots and stored in liquid nitrogen till use. In the text and in figures, Kinesin-1-GFP is represented in cerulean blue (False color LUT).

### Preparation of HEK293 cell lysates

0.6 million HEK293 cells/well were seeded into two 6-well plates. After 48 h, cell lysates were prepared according to the protocol mentioned in [56]. Briefly, transfected cells were treated with trypsin and centrifuged at 450xg for 10 min at 4 °C. All steps after centrifugation was done on ice or at 4°C. The cell pellet (from two 6 well plates) was resuspended in 130 µl ice-cold lysis buffer ((BRB80 containing 0.05% Triton X-100 and protease inhibitors (10 µg ml^−1^ leupeptin, aprotinin and 4-(2-aminoethyl)-benzenesulfonyl fluoride; Sigma-Aldrich)). The resuspended mixture was transferred to an ice-cold 1.5 ml Beckman ultracentrifuge tube, and the cells were further lysed by pipetting. This was followed by sonication (4 short pulses at 12% Amplitude, MS-72 probe, Bandelin Sonoplus). The lysed mixture was further mixed by pipetting and then centrifuged at 33,800xg for 30 min at 4°C. The supernatant was aliquoted, snap-frozen in liquid nitrogen and stored at −80 °C. The total protein concentration was estimated using Bradford method (Pierce^TM^ Bradford protein assay kit, 23200, ThermoScientific). For buckling assays, 20 µg ml^-1^ of HEK293 cell lysate (final concentration) was added along with ATP buffer.

### Preparation of GMPCPP stabilized microtubule seeds

Microtubule seeds were prepared by polymerizing 10 µM of tubulin (40% red fluorescent tubulin and 60% unlabeled or biotinylated tubulin) in BRB80 buffer supplemented with 0.5 mM GMPCPP (Guanosine-5’-[(α,β)-methyleno] triphosphate: slowly hydrolysable GTP analogue) (NU-405, Jena Bioscience) for 1 hour at 37°C. Following this, the above mixture was incubated with 1 µM Paclitaxel (Sigma) for 30 min at room temperature, centrifuged (21,300xg at 25°C for 15 min) and the pellet was resuspended in warm BRB80 supplemented with 0.5 mM GMPCPP. The prepared microtubule seeds were stored in liquid nitrogen until use.

### Preparation of Capped GDP microtubules for *in vitro* assays

Capped GDP microtubules in a single color (displayed in figures as agenta) were prepared with a lower proportion of fluorescent tubulin (12%) in the GDP lattice in contrast to the stabilized ends (40% labeled seeds and caps). Capped GDP microtubules were prepared by elongating prepared microtubules seeds with 10 μM of tubulin (12% labeled) in Elongation buffer (56 mM PIPES, 0.7 mM EGTA, 0.7 mM Mgcl_2_, 38 mM KCl, 19 mM Phosphate buffer, pH 6.8), supplemented with 1 mM GTP for 40 min at 37°C. The sample with the polymerized microtubules was then immediately centrifuged (21,300x g for 15 min). The resulting pellet was resuspended in a resuspending mix containing Elongation Buffer supplemented with 0.5 mM GMPCPP and further incubated at 37°C to cap the ends of the microtubule. Capping helps prolong the lifetime of the microtubule and additionally allows for better visualization of lattice self-repair. The resuspended mix is then successively capped by adding 0.5 μM of tubulin (60% biotin, 40% fluorescent tubulin) in a stepwise manner (for a total of 10 times) every 15 min.

For microtubule static curvature and double anchored motor assays, GDP microtubules with seeds and caps containing biotin were prepared. For microtubules buckling assays, GDP microtubules were prepared using biotin-containing seeds and the caps were grown using 0.5 µM (60% unlabeled and 40% fluorescent) tubulin. For microtubules gliding assays, GDP microtubules were prepared using seeds (60% unlabeled and 40% fluorescent, without biotin) and the caps were grown using 0.5 µM tubulin (60% unlabeled and 40% fluorescent).

### Cover-glass passivation

For experiments analyzing static curvature of microtubules, cover-glasses were passivated with Silane-PEG-Biotin as follows: Cover-glasses were first wiped with lint-free KimWipe^TM^ (Kimberly-Clark Professional™ 33670-04) tissues and 70% ethanol, incubated in Acetone for 30 min, followed by 96% ethanol for 15 min (gentle agitation at room temperature). The cover-glasses were subsequently rinsed in ultrapure water, incubated in Hellmanex III solution (2% in water, Hellmanex) for 2 h (gentle agitation at room temperature), washed in ultrapure water and dried. This was followed by treatment using an Deep-UVO cleaner (30 mW/cm² at 254 nm, 144AX-220 Jelight) for 30 min before incubation in 7:3 mix of tri-ethoxy-silane-PEG and tri-ethoxy-silane-PEG-biotin (30kDa, Creative PEG works)(final concentration of 1 mg ml^-1^ in 96% ethanol and 0.1% HCl), for 3 days with gentle agitation at room temperature. After 3 days, the PEGylated cover-glasses were extensively rinsed in 96% ethanol and ultrapure water, air-dried and stored at 4°C till use.

For gliding and buckling assays with kinesin-1 motor proteins, cover-glasses were wiped with lint-free KimWipe^TM^ tissues and 96% ethanol, rinsed in ultrapure water and sonicated in 2% Hellmanex-III solution at 60°C for 30 min. Following sonication, the cover-glasses were rinsed in ultrapure water, stored in ultrapure water at room temperature and air-dried just before use.

### Assay to assess self-repair in static bent microtubules

A flow chamber of an approximate volume of 60 μl was built by sandwiching two pieces of SiPEG-Biotin passivated cover-glasses using two strips of double-sided adhesive tape (70 μm height, 0000P70PC3003, LiMA, Couzeix, France). The flow chamber was first perfused with streptavidin (100 μg ml^-1^ in 1xBRB80; Fisher Scientific) for 1 minute. This was followed by a solution of 0.1 μg ml^-1^ PLL-g-PEG (Pll 20 K-G35-PEG2K, Jenkam Technology, in 10 mM Na-HEPES, pH 7.4) for 1 minute. Following a wash step with 1xBRB80, prepared capped GDP microtubules (diluted in 1xBRB80) were flushed in an alternative manner (to achieve bending) and incubated for 5 min before extensive washing with BRB80 supplemented with 1 mg ml^-1^ BSA (Bovine Serum Albumin, Sigma). Subsequently, an incorporation mix containing 5 µM Tubulin (100% ATTO-488, green labeled) in Elongation buffer supplemented with 1 mM GTP, an oxygen scavenger cocktail (22 mM DTT, 1.2 mg ml^−1^ glucose, 8 μg ml^−1^ catalase and 40 μg ml^−1^ glucose oxidase), 1 mg ml^−1^ BSA and 0.033% (w/v) methyl cellulose (1,500 cP, Sigma) was added and allowed to incubate at 37°C for 15 min. Following incubation, the chamber was perfused with an imaging buffer (composition same as that of the incorporation buffer with an addition of 2 µM unlabeled tubulin-to keep microtubules stable for imaging). The chamber was then sealed with Valap before imaging.

### Microtubule gliding assay

*In vitro* gliding assays were performed in 20 μl flow chambers constructed from Hellmanex-sonicated cover-glasses and using double-sided tape. The chamber was first perfused with a 20 μl solution of 0.2 mg ml^-1^ Anti-Green-Fluorescent-Protein (GFP) Antibody (Invitrogen, A-11122) for 3 min. This was followed by further passivation using 1% w/v BSA solution in 1xHKEM buffer (10 mM HEPES buffer (pH 7.2), 50 mM KCl, 1 mM EGTA and 5 mM MgCl_2_). Kinesin-1-GFP diluted in TicTac buffer (10 mM HEPES buffer (pH 7.2), 16 mM PIPES buffer (pH 6.8), 50 mM KCl, 5 mM MgCl_2_, 1 mM EGTA, 20 mM dithiothreitol (DTT), 3 mg ml^−1^ glucose, 20 µg ml^−1^ catalase, 100 µg ml^−1^ glucose oxidase and 0.3% BSA) was added and allowed to incubate for 5 min. Subsequently the chamber was washed with TicTac buffer and prepared capped GDP microtubules (diluted in 1xBRB80) was added and allowed to incubate for 2 min, followed by extensive washing with TicTac buffer. ATP Buffer (10 mM HEPES buffer (pH 7.2), 56 mM PIPES buffer (pH 6.8), 50 mM KCl, 5 mM MgCl_2_, 1 mM EGTA, 20 mM dithiothreitol (DTT), 3 mg ml^−1^ glucose, 20 µg ml^−1^ catalase, 100 µg ml^−1^ glucose oxidase, 0.3% BSA supplemented with 2.7 mM of ATP and 0.2% methyl cellulose) was then added to initiate motor activity. The chamber was sealed with Valap and immediately imaged. For control experiments, ATP was replaced with 2.7 mM of AMPPNP (Adenosine-5′-(β,γ-imido) triphosphate; A2647, Sigma-Aldrich) in the final buffer mix. For experiments with survival in the presence of free tubulin, 5 µM tubulin (100% unlabeled) was added to the ATP buffer. For gliding assays in the presence of HEK293 cell lysate, 20 µg ml^-1^ of cell lysate was added in the ATP buffer (all other steps were kept the same).

### Microtubule buckling assay

To achieve microtubule buckling, the flow chamber was first perfused with 100 μg ml^-1^ of streptavidin (diluted in 1xHKEM, Invitrogen, 434301) prior to the anti-GFP-antibody step. Capped GDP microtubules made from biotinylated seeds were used (all other steps were kept the same). All other steps were the same as the microtubule gliding assay described above. For buckling assays in the presence of HEK293 cell lysate, 20 µg ml^-1^ of cell lysate was added in the ATP buffer.

### Incorporation in microtubule gliding and buckling assays

To visualize self-repair (incorporation of free tubulin dimers) in buckling and gliding microtubules, an additional coating step (TicTac buffer supplemented with 2 μM unlabeled tubulin) was performed prior to addition of prepared GDP microtubules as free green tubulin dimers tend to attach to the layer of motors during the assay, effectively hindering the detection of the incorporated dimers [9]. To visualize self-repair, gliding and buckling microtubules were exposed to ATP buffer supplemented with 5 μM tubulin (100% labeled) and allowed to glide/buckle at 37°C for 15 min. Following incubation, the chamber was perfused with ATP buffer containing 2 µM unlabeled tubulin (to stabilize microtubules for imaging), sealed with Valap before imaging.

### Single molecule photobleaching experiments to determine kinesin surface density

Single molecule photobleaching (SMPB) was performed according to the protocol described previously in [57]. Briefly, a 1µM dilution of Kinesin-1-GFP in cold 1xHKEM was centrifuged to remove aggregates (15 min, 4°C, 215,000x g in a Type 70 Ti rotor [Beckman Optima XPN80]) prior to the assay. In a 20µl flow chamber made from Hellmanex sonicated cover-glasses, 20 µl of 0.2 mg ml^-1^ solution of Anti-GFP antibody was flushed in and incubated for 3 min. This was followed by a 1xHKEM wash step and then a 350 pM solution of Kinesin-1-GFP (further diluted in 1xHKEM) was added and incubated for 5 min. The chamber was then washed with 300 µl of 1xHKEM to remove unbound Kinesin-1 molecules and sealed using Valap. Photobleaching was achieved at 65% Laser power with 500 ms exposure time (for 5 min) in continuous streaming mode. Recorded time-lapses were cropped and regions of interest (ROIs) with uniform illumination were chosen for analysis. Using Stowers Institute Fiji plugin, the fluorescent intensity traces of individual Kinesin-1-GFP molecules (represented in cerulean blue in **ED Fig. 4a(i)**) were obtained. From the step-like traces (refer **ED Fig. 4a(ii)**), we quantified the average fluorescent intensity of one Kinesin-1 molecule. To quantify the surface density of Kinesin-1 (refer **ED Fig. 4b**) at our working concentration of 416 nM, we repeated the above assay with 416 nM of kinesin-1-GFP and captured images using the same imaging conditions (65% Laser power with 500 ms exposure time, same camera gain and binning settings as above). ROIs with uniform illumination were chosen for analysis and average intensity of fluorescent Kinesin-1 at 416 nM was estimated. The experiment was performed on the same day and using the same imaging settings as the SMPB assay. The surface density (No: of Kinesins/µm^2^) was estimated by dividing the average intensity of 416 nM Kinesin/ [(Average intensity of one Kinesin-1 molecule) * (Pixel size)^2^].

### Single-Molecule motility experiments to estimate kinesin motility parameters

For single molecule experiments to estimate motility parameters of Kinesin-1-GFP molecules (represented in cerulean blue in **ED Fig. 4**), an orbital TIRF microscope was used. Prior to the assay, a 1 µM dilution of Kinesin-1-GFP in cold 1xHKEM was centrifuged to remove aggregates (15 min, 4°C, 215,000x g). In a 20 µl flow chamber made from PEGylated cover-glasses, 20 µl of streptavidin (50 µg ml^-1^ in 1xHKEM; Fisher Scientific) was flushed in and allowed to incubate for 1 min. Following a wash step with 1xHKEM, prepared capped GDP microtubules with biotinylated seeds and caps (diluted in 1xBRB80) were flushed in and incubated for 5 min before extensive washing with 1xHKEM. Subsequently, motility buffer containing 500 pM Kinesin-1-GFP in TicTac buffer (containing the oxygen scavenger solutions to minimize bleaching of fluorophores) was flushed in. One still image of the microtubule was acquired. Motile kinesin-1 single molecules were captured in continuous streaming mode in the GFP channel (using the same image conditions as in the SMPB assay). The time-lapses were processed in Fiji and kymographs (Refer **ED Fig. 4c**) were generated using the KymoResliceWide plugin. Tracking the traces of kinesin-1-GFP molecules from kymographs from 50 frames with a temporal cutoff of 500 ms, distributions of the run length (distance traveled by an individual kinesin-1 molecule on a microtubule), dwell time (total residence time of individual kinesin-1 molecule on a microtubule), mean velocity and detachment rate of kinesin-1 molecules were estimated (**ED Fig. 4d-g)**.

### Imaging

63x oil immersion objective (Zeiss, Plan-Apochromat 63x/1.40 oil DIC M27) of a Zeiss LSM 900 confocal microscope with an Axiocam 705 Mono camera (Zeiss) was used for imaging. The microscope stage was kept at 37°C by means of a warm stage controller (Insert-P, PeCon). The temperature on the microscope stage was controlled with the incubator (PeCon) kept at 37°C. Time-lapses were recorded using ZenBlue software (version 3.2, Zeiss). Single molecule experiments were performed on an objective-based orbital TIRF microscope (Nikon Ti2 Eclipse, modified by ViSitron Systems) equipped with an EMCCD Camera (Andor iXon Life). The microscope stage was maintained at 37°C using a warm stage controller (OkoLabs). Time-lapses were recorded using VisiView software (version 6.0). Microtubule thermal fluctuations time-lapses were imaged using the 60x Oil immersion objective of a Nikon Ti2E Epifluorescence microscope using NIS elements AR software (Nikon).

### Image processing and quantification of incorporation stretches

Videos were processed to improve the signal/noise ratio (subtract background and smooth functions of Fiji, version 1.53t) [58]. Self-repair events or incorporations were estimated from overlaid images (average of 3 frames) from time-lapses taken every 2 s. Incorporation stretches were identified from line scans (green fluorescence) along the magenta GDP microtubule lattice. A stretch of green fluorescence along the microtubule (in magenta as indicated in **Fig. 1c**) was regarded as an incorporation if it displayed a fluorescence intensity higher than 1.5 times that of the background as well as followed the lateral fluctuations of the microtubule. The full-width-half-maximum (FWHM) distance from the intensity profile of the incorporation stretch was taken as the incorporation length. Accounting for the resolution limit of the microscope, incorporation stretches spanning less than 250 nm were disregarded from analysis.

For data on microtubule self-repair in cells, images of a microinjected cell were first divided into 4-5 ROIs (regions of interest, as indicated in **Fig. 1g** with a box with a red outline) and sections of microtubules (indicated with boxes with white outline in **Fig. 1g**) were analyzed. Incorporations found on bundled microtubules and at microtubule crossover sites (and up to 0.8 μm away from crossover sites) were disregarded from analysis. As mentioned in [29], longer incorporation stretches were found in microtubules located close to the microinjection sites owing to possible damage from microinjection. Hence, only microtubule sections found 10 μm or more from the microinjection site were considered for analysis.

### Curvature analysis

Images of both straight and bent microtubules were tracked using the Fiji plugin Jfilament2D [59] and the curvature of microtubules were estimated using a custom-made python script (see **ED Fig. 2** for analysis workflow). The obtained curve from JFilament2D was first smoothed by a parametric spline interpolation to remove noise. Then, the Menger curvature was calculated using three points spaced 120 nm apart. Curved microtubules displaying a maximum local curvature of 0.15 µm^-1^ or above were considered as bent. Bent zones of bent microtubules refer to sections of the bent microtubule with a mean local curvature above 0.2 µm^-1^. Mean curvature over the entire microtubule was estimated by averaging the local curvature of all segments of the traced microtubule. Curvature at point of breakage of breaking buckling microtubules was estimated by taking an average of the local curvature of all segments in the region spanning 500 nm around the point of breakage.

### Estimation of the microtubule persistence length

For thermal fluctuation experiments to estimate the microtubule persistence length, microtubules were elongated from biotin containing microtubule seeds (attached to a streptavidin coated SiPEG-biotin cover-glass) using elongation buffer supplemented with 1 mM GTP and 11 µM tubulin for 15 min at 37°C. Following this the elongated microtubules were capped by flushing in a solution of elongation buffer supplemented with 0.5 mM GMPCPP and 2.5 µM tubulin and allowed to incubate at 37°C for 5 min. After washing with imaging buffer, the chamber was sealed and thermal fluctuations of microtubules were captured every 10 s for 40 min. Images were analyzed in Fiji and the filament coordinates were obtained using T-SOAX software [60] and the persistence length of each microtubule was estimated following the method reported in [25]. Using a custom-written python code, the tangent angles were obtained via finite differences and expressed as cosine modes. The persistence length was obtained by fitting variance of the cosine modes (Refer **ED Fig. 4i**).

### Quantification of lateral tubulin incorporation and % of tubulin turnover

The amount of lateral tubulin incorporation in incorporated stretches in all datasets was estimated by using stretches of microtubule elongation (stretches of 100% green-labeled tubulin that we occasionally observed beyond the cap-Refer **Fig. 1j, top**) as a reference stretch that we assumed to possess a 13-protofilament structure. As described in [5,8], using the estimates of the length (FWHM), the integral (I_total_) fluorescence intensity of the reference stretch, length of the tubulin dimer (L= 8 nm), we estimated the fluorescence intensity of a single tubulin dimer as (I_dimer_) as I_dimer_=I_total_*L /(FWHM*13). From the integral fluorescence intensity (I_inc_) of the incorporation spot, we estimated the number of incorporated dimers as N_inc_=I_inc_/I_dimer_. The values of N_inc_ from five frames per incorporation stretch were averaged to obtain the amount of lateral tubulin incorporation per incorporation in each condition. To estimate the tubulin turnover % in all conditions, we multiplied the mean value of the incorporation length/microtubule length (estimate of the lattice length replaced) with the average amount of lateral incorporation in each condition.

### Categorization of different microtubule bending events in cells

Time-lapses of dynamic microtubules in live PtK2 cells were analyzed for quantification of microtubule bending events in cells. The bending events in cells were classified in to ‘Bent persisting’, ‘Buckling’, ‘Looping’ and ‘Breakage’ events (See **ED Fig. 3a**). A microtubule was considered to persist in the bent form without relative change in its curvature (less than 0.1-0.15 µm^-1^) (like in **Fig 2b(i)**) for the period of observation (5 min). Microtubules showing dynamic change in curvature (See **Fig. 2a**, **2b(ii)**) were classified as buckling. Buckling microtubules that were seen to adopt a loop-like conformation were categorized as looping microtubules. We also observed relatively rare instances of microtubule breakage following bending/buckling (like in **Fig. 2b (iii)** and **ED Fig. 3b**). A total of 4 cells from two independent experiments were analyzed.

### Microtubule survival

For microtubule survival experiments, time-lapses were recorded for 40 min with a frame interval of 10 seconds. The % microtubule population remaining was estimated by manually counting the no: of microtubules in each frame within the same field-of-view every 2 min.

### Estimation of kinesin motility parameters

Gliding velocity of microtubules both in the presence and absence of HEK293 cell lysate was estimated by tracking the movement of a gliding microtubule in 10 consecutive frames using MTrackJ Fiji plugin. Kinesin motility parameters were estimated from kymographs generated from traces using the KymoResliceWide plugin.

### Statistical analysis

Statistical analysis was performed using GraphPad Prism software (version 9.5). To test the significance in the case of incorporation lengths and amount of lateral tubulin incorporation, Mann-Whitney test (two-tailed) was used as a non-parametric alternative to a t-test, as the distributions are non-Gaussian in nature. For comparisons of incorporation frequency, incorporation length/microtubule length, unpaired-t-test was used as the distributions have similar variances and are Gaussian in nature.

## Contributing authors (alphabetical order)

Albrecht, Constantin Matteo

Bosche, Jonas

Diez, Stefan

Grünewald, Mona

König, Belinda

Nandakumar, Shweta

Santen, Ludger

<u>Schaedel, Laura</u>

<u>Shaebani, Reza</u>

Wieczorek, Mirko

LaS, RS, SD, LuS conceptualized and guided the project. LaS, SN, SD and MW designed the experiments. SN carried out the experiments and analysis. MG and BK purified the tubulin. MW assisted with image and curvature analysis. CMA assisted with cell lysate experiments and design of figure schematics. RS, LuS and JB designed, performed and analysed the simulations. LaS and RS wrote the manuscript. All authors provided critical feedback.

## Funding sources

LuS, LaS, SD and RS were supported by the DFG grant SFB 1027. The work was financially supported by the European Research Council (ERC; grant no. StG 101115795 to LaS).

The authors are thankful to Jérémie Gaillard and Laurent Blanchoin (Cytomorpho Lab, Grenoble) for sharing reagents and technical expertise. We are grateful to Morgan Gazzola and Manuel Théry (Cytomorpho Lab, IPGG, Paris) for sharing their previously published dataset on microtubule self-repair in cells. We thank Cordula Reuther and Rahul Grover (B-CUBE, TU Dresden) for sharing kinesin related expertise, Subham Biswas (Experimental Physics and Center for Biophysics, University of Saarland) for assistance with single molecule imaging and Karin John (LIPhy, Université Grenoble Alpes) for fruitful discussions.

## Ethics declaration

The authors declare no competing interests.

## Data availability

Curvature analysis code and source data underlying the main and supplementary figure plots will be made available in the Zenodo repository link: 10.5281/zenodo.16935847 upon publication. The custom simulation code is available upon reasonable request by contacting the corresponding authors.

---

**Extended Data Table 1:** Parameters used in numerical simulations of microtubule deformation under active forces. For experimental data values obtained from experiments, please refer ED Fig 4.

| Parameter | Symbol | Value | Unit | Reference |
| --- | --- | --- | --- | --- |
| Thermal persistence length | $L_p$ | [1,10] | mm | This study (ED Fig 4i) |
| Default thermal persistence length | $L_{p,0}$ | 5 | mm | This study (ED Fig 4i) |
| Microtubule length | $L$ | [5,15] | $\mu\text{m}$ | This study |
| Default microtubule length | $L_0$ | 10 | $\mu\text{m}$ | This study (ED Fig 4h) |
| Microtubule discrete units | $\Delta L$ | 8 | nm | [39],[40] |
| Hopping step size | $d_h$ | 8 | nm | [39],[40] |
| Attachment rate | $\omega_a$ | [1,10] | $\text{s}^{-1}$ | [40],[61],[62] |
| Default attachment rate | $\omega_{a,0}$ | 5 | $\text{s}^{-1}$ | [40],[61],[62] |
| Detachment rate | $\omega_{d,0}$ | 0.5 | $\text{s}^{-1}$ | This study (ED Fig 4g), [61], [62] |
| Hopping rate | $\omega_{h,0}$ | 81 | $\text{s}^{-1}$ | [61] |
| Motor density | $\rho$ | [100,500] | $\mu\text{m}^{-2}$ | This study |
| Default motor density | $\rho_0$ | 400 | $\mu\text{m}^{-2}$ | This study (ED Fig 4b) |
| Spring constant | $k$ | 0.0003 | N/m | [63] |
| Detachment force | $f_d$ | 3 | pN | [64],[65] |
| Attachment force | $f_a$ | 3 | pN | [66],[67] |
| Motor stall force | $f_s$ | 6 | pN | [68],[69] |

**Extended Data Figure 1:**
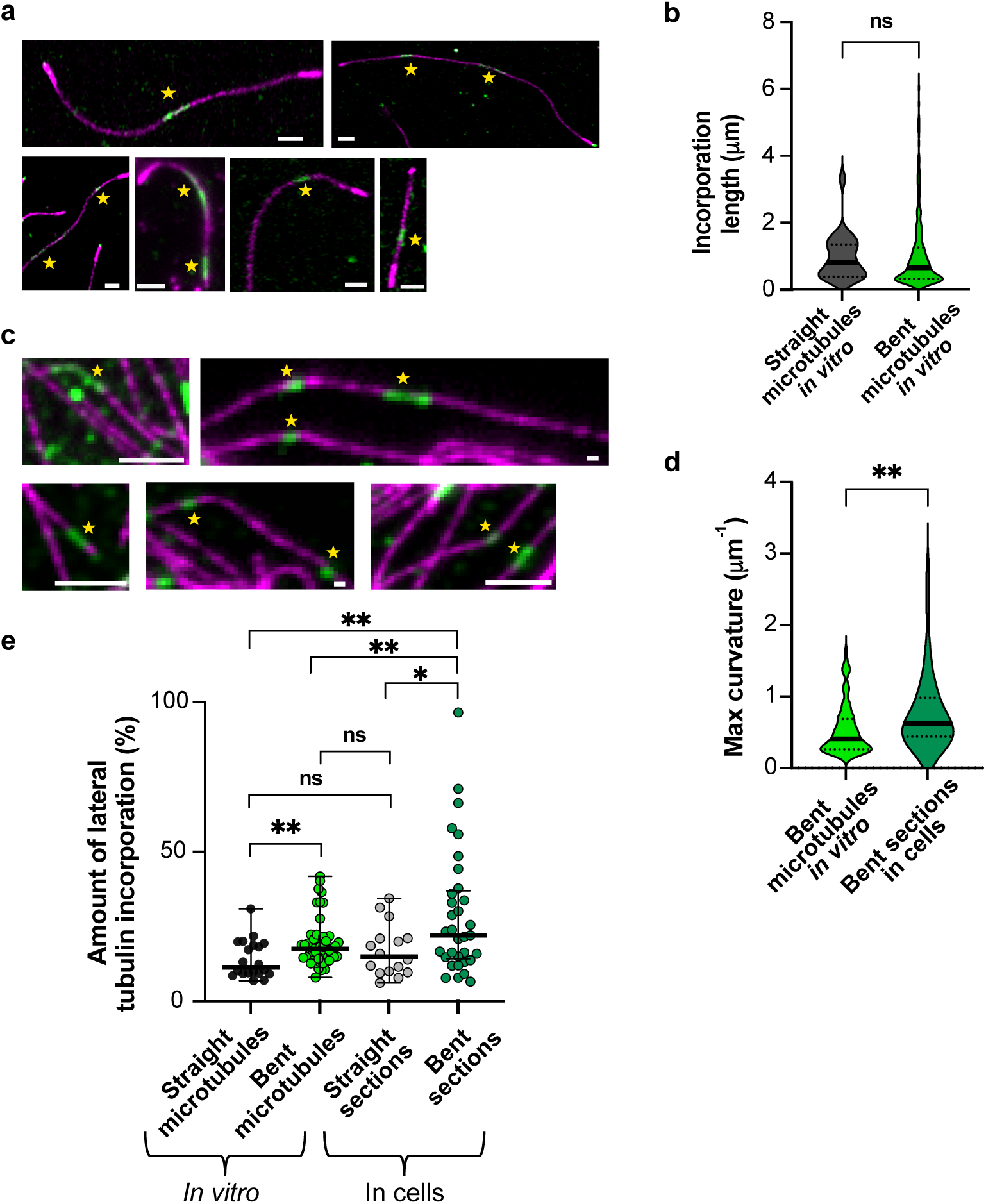
**a**, Additional examples of incorporations (marked with a yellow star) in static straight and bent microtubules *in vitro* (scale bar: 2 µm) **b**, Comparison of incorporation length in static straight vs static bent microtubules *in vitro* (global). Black line represents the median and dotted lines, the interquartile range. p= 0.2905 (ns, not significant) using Mann-Whitney test. Total length of microtubule analyzed in both cases: 1,104 µm (n= 23 incorporations for straight microtubules and n= 263 incorporations for bent microtubules) **c**, Additional examples of incorporations (marked with a yellow star) in straight and bent sections in cells (scale bar: 2 µm) **d**, Comparison of curvature of static bent microtubules *in vitro* vs bent sections in cells. Black line represents the median and dotted lines, the interquartile range. p= 0.027 using an unpaired t-test (n= 127 *in vitro*; n= 36 in cells). **e,** Higher amount of lateral tubulin incorporation in bent microtubules, both in cells and *in vitro*. Three independent experiments were analyzed for each condition. n= 21 incorporations for straight microtubules (*in vitro)*, n= 51 incorporations for bent microtubules (*in vitro)*, n= 16 incorporations for straight sections (cells) and n= 32 incorporations for bent sections (cells). Black lines represent the median and error bars represent the interquartile range. Statistical test used: Mann-Whitney test. p= 0.0391 (straight sections, cells -bent sections, cells); p= 0.0024 (straight microtubules, *in vitro*-bent microtubules, *in vitro*); p= 0.0095 (bent sections, cells-bent microtubules, *in vitro*); p= 0.2411 (not significant; straight sections, cells-straight microtubules*, in vitro*); p= 0.2163 (not significant; straight sections, cells-bent microtubules, *in vitro*) and p= 0.2411 (not significant; straight sections, cells-straight microtubules, *in vitro*).

**Extended Data Figure 2:**
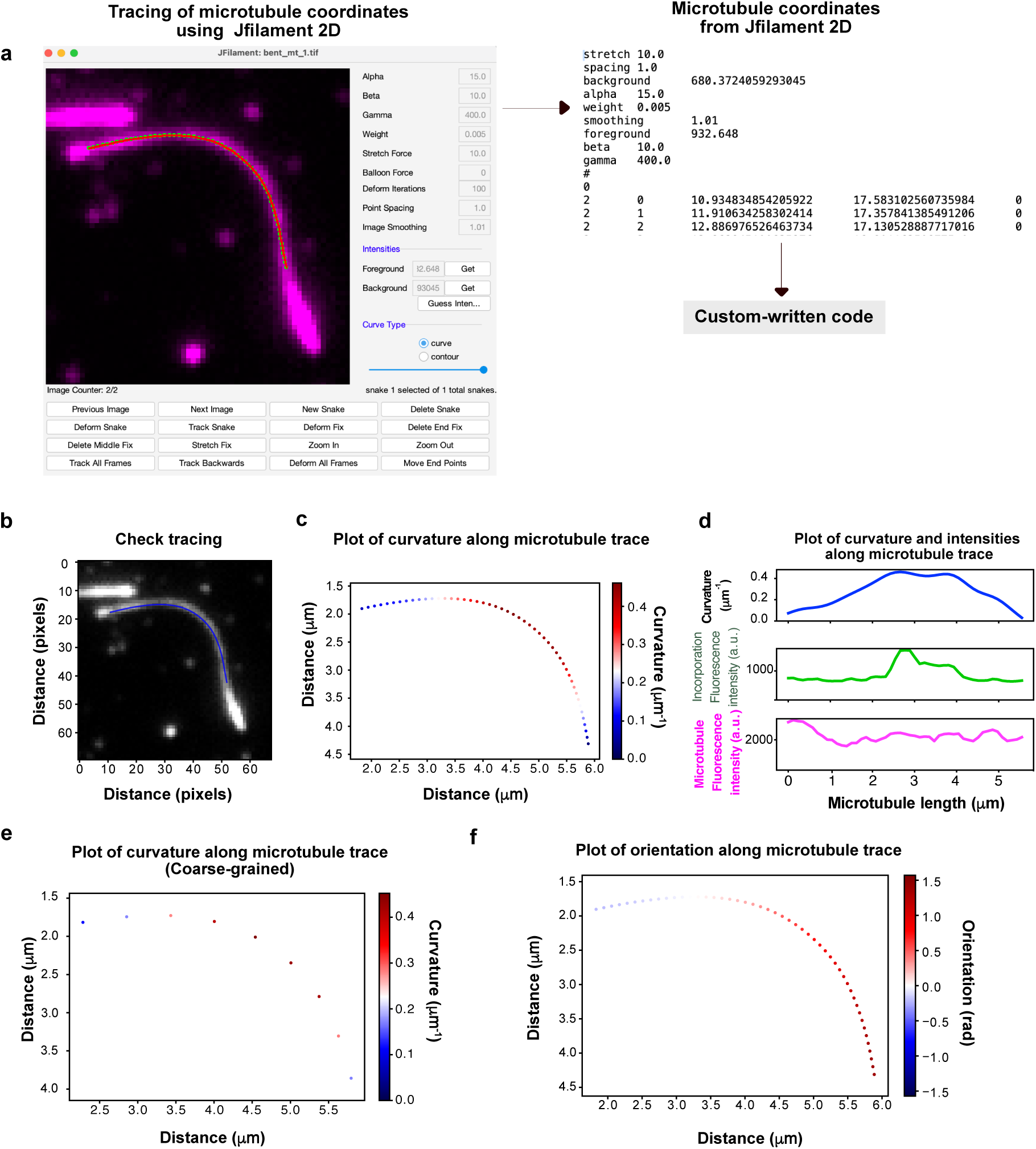
Curvature analysis workflow. **a,** Microtubules were traced using the Fiji plugin JFilament 2D. The obtained coordinates were used to calculate the curvature and to plot the fluorescence intensities along the microtubule arclength using a custom-written code. **b,** *Output 1 from code:* Image of traced microtubule superimposed on the original microtubule image to check for accuracy of tracing. Checking the accuracy of tracing might be necessary if the obtained curve from JFilament 2D was additionally smoothed in the code. **c,** *Output 2 from code:* Plot of curvature (color-coded) along the traced microtubule **d,** *Output 3 from code:* Plot of curvature (µm^-1^) and fluorescence intensities (a.u.) of both microtubule and incorporation channel along the microtubule length (µm). **e,** *Optional output 4 from code:* Plot of coarse-grained curvature (color-coded) along the microtubule segments. **f,** *Optional output 5 from code:* Plot of orientation (color-coded) along the traced microtubule. Microtubules showing a local curvature higher than 0.15 μm^-1^ were considered as bent.

**Extended Data Figure 3:**
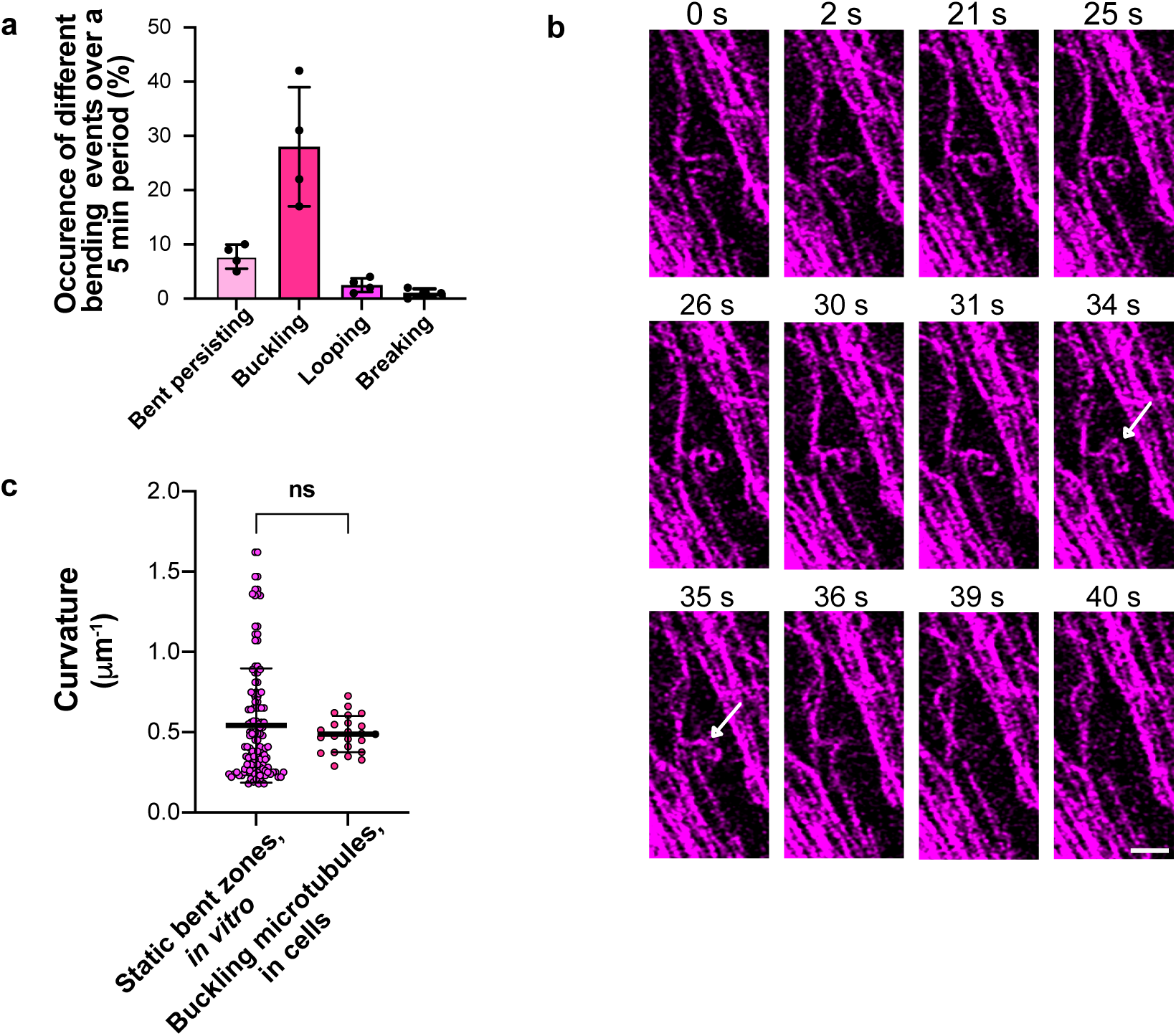
**a,** *Quantification of different microtubule bending events* over a 5 min period in live PtK2 cells (with an endogenous tubulin-eGFP tag; represented in magenta). Error bars represent the S.D. Black dots represent individual cells. n= 4 cells analyzed from two independent experiments. **b,** Time-lapse sequence showing breakage at high curvatures (marked by white arrow) in a looping microtubule in a live PtK2 cell. Scale bar: 2 µm. **c,** Comparison of curvature between the bent zones of static bent microtubules (*in vitro* analyzed in Fig 1d) and buckling microtubules (in cells, analyzed in Fig 2g). Black line represents the mean and error bars represent the S.D. (n= 127 microtubules from static bent zones *in vitro* and n= 23 buckling microtubules, *in vitro* from three independent experiments). p= 0.4721 (not-significant; ns) using unpaired-t-test.

**Extended Data Figure 4:**
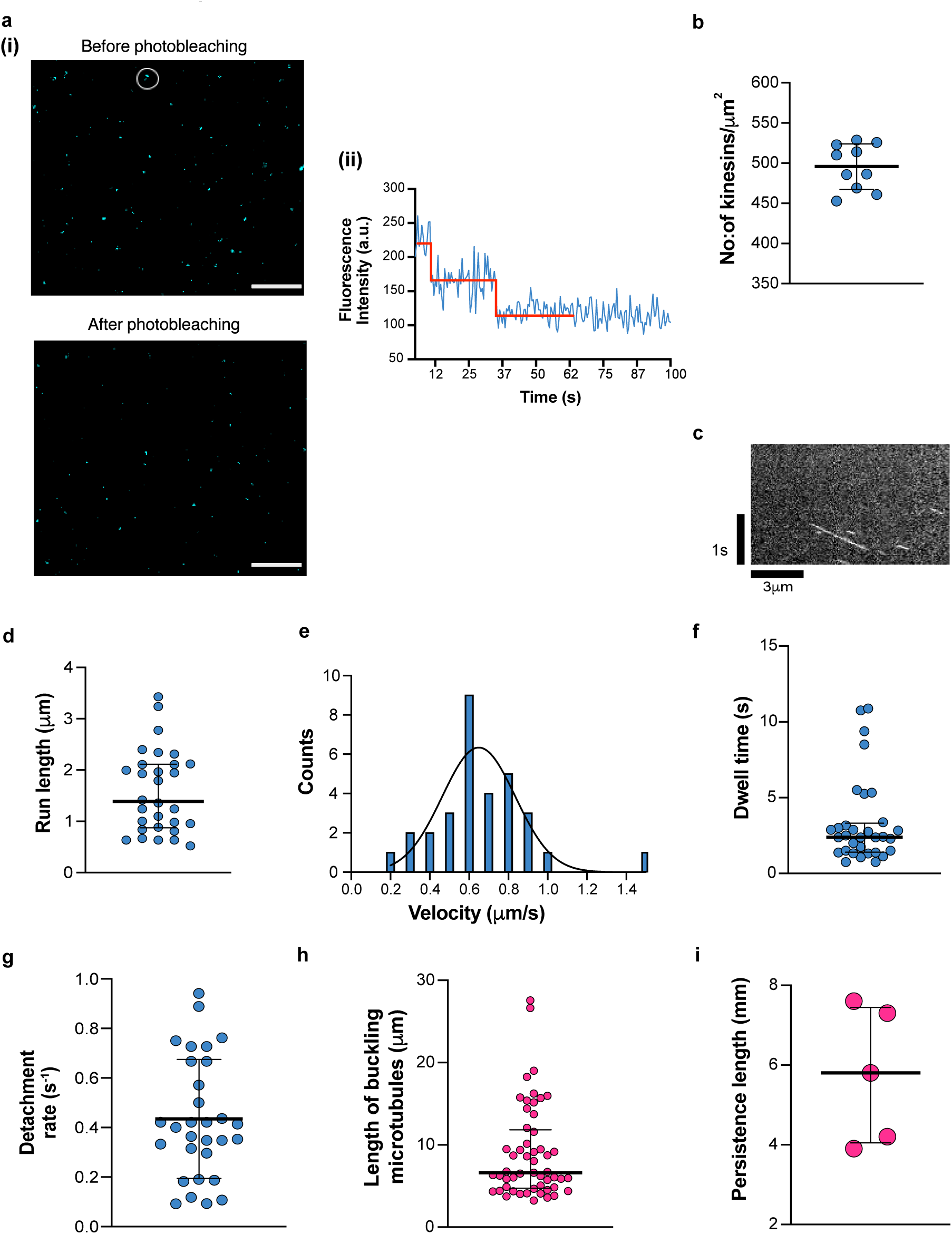
Data underlying experimental parameters for simulations: Single molecule photo-bleaching to estimate kinesin surface density: a,(i) Example image of kinesin-1-GFP (represented in cerulean blue) molecules at 350 pM before and after photobleaching using 65 % laser power in streaming mode, with 500 ms exposure time. White circle indicates a kinesin-1-GFP molecule that disappears over time due to photobleaching. Scale bar: 5 µm. a,(ii) Trace showing two-step photobleaching of the molecule circled in a(i). Red line is overlaid for better visualization of the two-step drop in fluorescence intensity b, Plot showing distribution of the surface density of kinesin-1 estimated from photobleaching experiments. Black line represents the mean and bars represent the S.D (n= 10 traces from 10 spots were analyzed from two independent experiments). *Single molecule motility studies to determine kinesin motility parameters:* c, Exemplary kymograph showing motile kinesin-1 molecules. d, Distribution of run length of kinesin-1 molecules. Black lines represent the median and the bars represent the interquartile range. e, Distribution of the velocities of kinesin-1 molecules. f, Distribution of dwell time of kinesin-1 molecules on microtubules. Black line represents the median and bars represent the interquartile range. g, Distribution of detachment rate of kinesin-1 molecules. Black lines represent the mean and bars represent the S.D. For the data underlying Fig ED 4d-g, n= 32 kymographs from 13 microtubules from two independent experiments were analyzed. h, Distribution showing length of buckling microtubules. Black lines represent the median and bars represent the interquartile range. (n= 53 microtubules from six independent experiments). i, Persistence length of microtubules from thermal fluctuation experiments. Black lines represent the median and bars represent the interquartile range (n= 5 microtubules from three independent experiments).

**Extended Data Figure 5:**
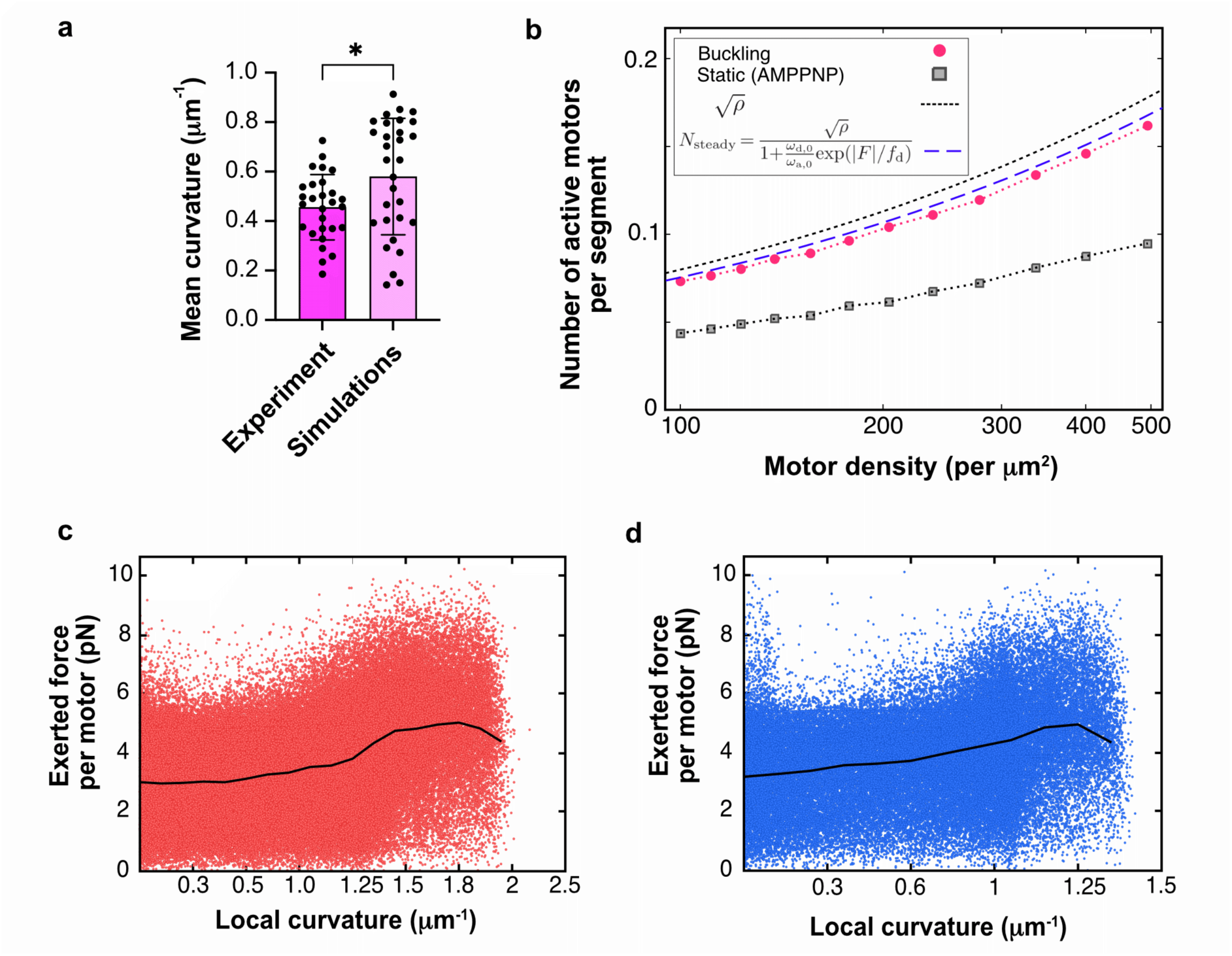
**a,** *Comparison of mean curvature of buckling microtubules in experiments vs simulations*. Simulation results using a motor density of ρ = 494 μm^−2^, microtubule persistence length, L_p_= 6 mm, and microtubule length, L= 10 μm. Error bars represent the S.D. p= 0.0174 using unpaired t-test (n = 28 microtubules from experiments and n= 30 microtubules from simulations). **b**, *Lin-log plot of number of active motors per segment in a simulated buckling microtubule as a function of motor density.* The steady-state number of motors bound per microtubule segment can be estimated by analyzing the Markov chain balance of stochastic binding and unbinding events, yielding a value for N_steady_ (see SI simulation methods for details). In this equation, |F| is the absolute value of the average force exerted on the filament by the motors (for both buckling and AMPPNP conditions) at different motor densities (See Fig 6i and ED Fig 6c). In practice, the actual number of motors on the filament is slightly lower than this theoretical estimate, as the system remains in a transient dynamical regime and motor traffic along the filament has not yet reached equilibrium. *Scatter plots of the force exerted by each motor vs local curvature*, shown for a: **c**, soft microtubule with L_p_= 1 mm (red dots) and **d**, stiff microtubule with L_p_ = 10 mm (blue dots). Other parameters include: L= 5 μm, ρ = 400 μm^−2^. Black line represents average values computed by binning the local curvature. Each plot includes data from 10^5^ microtubule samples. **Note:** Although the motor force values in the above scatter plots (ED 5c and 5d) span a broad range (even beyond 10 pN), most data points cluster at lower forces. The observed high forces occur due to the large ensemble size in simulations (10^5^ microtubule samples with ∼10^2^ motors along each). This enables us to observe statistically rare outcomes that would be difficult to capture in typical experimental datasets.

**Extended Data Figure 6:**
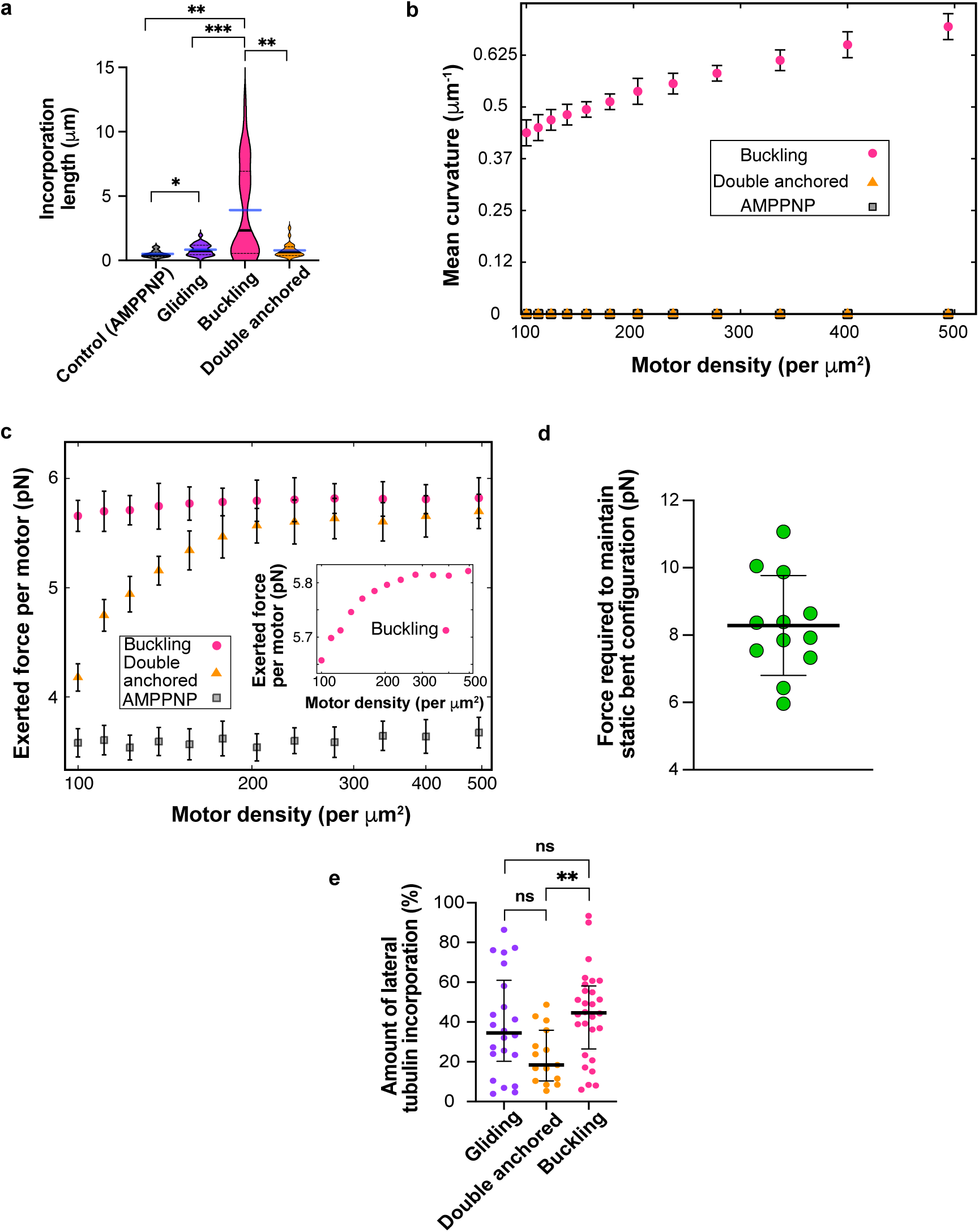
**a,** *Incorporation length across different conditions*-control (AMPPNP), gliding, buckling and double-anchored microtubules. Black lines represent median, dotted lines represent the interquartile range and blue line represents the mean from three independent experiments in each case, comprising 98, 139 and 110 μm of the total microtubule length analyzed for double-anchored microtubules. For microtubule lengths analyzed in all other conditions, refer Fig 3g captions. p= 0.3442 (double anchored-gliding); p= 0.0012 (double-anchored -buckling); p = 0.0018 (buckling-control), p= 0.0172 (gliding-control), and p= 0.0005 (gliding-buckling) using Mann-Whitney test. Comparison between all other conditions are non-significant. **b,** *Effect of motor arrangement and mobility on microtubule dynamics*. Simulations compare three configurations: (pink circles) motors distributed across the surface leading to buckling, (orange triangles) motors arranged linearly beneath an straight microtubule (double-anchored), and (gray squares) in the presence of AMPPNP. Mean curvature of each discretized node of the microtubule is plotted as a function of motor density for all three cases. Microtubule parameters: L= 10 μm and L_p_ = 5 mm. Data are averaged over time and across five microtubules. **c,** *Mean force exerted per motor on a microtubule as a function of motor density in simulations* for buckling, double anchored and AMPPNP conditions. Microtubule parameters: L= 10 μm and L_p_= 5 mm. The inset shows the same data as the buckling case in the main figure panel in the same figure, but over a narrower force range to better highlight the increasing trend. **d,** *Force (along the entire microtubule) required to maintain static bent microtubules in the bent configuration*. Refer to SI simulation methods for details. Black lines represent mean and S.D. (n= 12 static bent microtubules were analyzed from experimental data from three independent experiments). **e,** Higher amount of lateral tubulin incorporation in buckling microtubules. Black lines represent the median and bars represent the interquartile range. Statistical test used: Mann-Whitney test. p= 0.42, not significant (gliding-buckling); p= 0.1049, not significant (gliding-double-anchored); p= 0.0019 (double-anchored-buckling). 98, 139 and 110 μm of the total microtubule length analyzed for double-anchored microtubules. For microtubule lengths analyzed in all other conditions, refer Fig 3g captions. Data cumulative for three independent experiments in each condition.

**Extended Data Figure 7:**
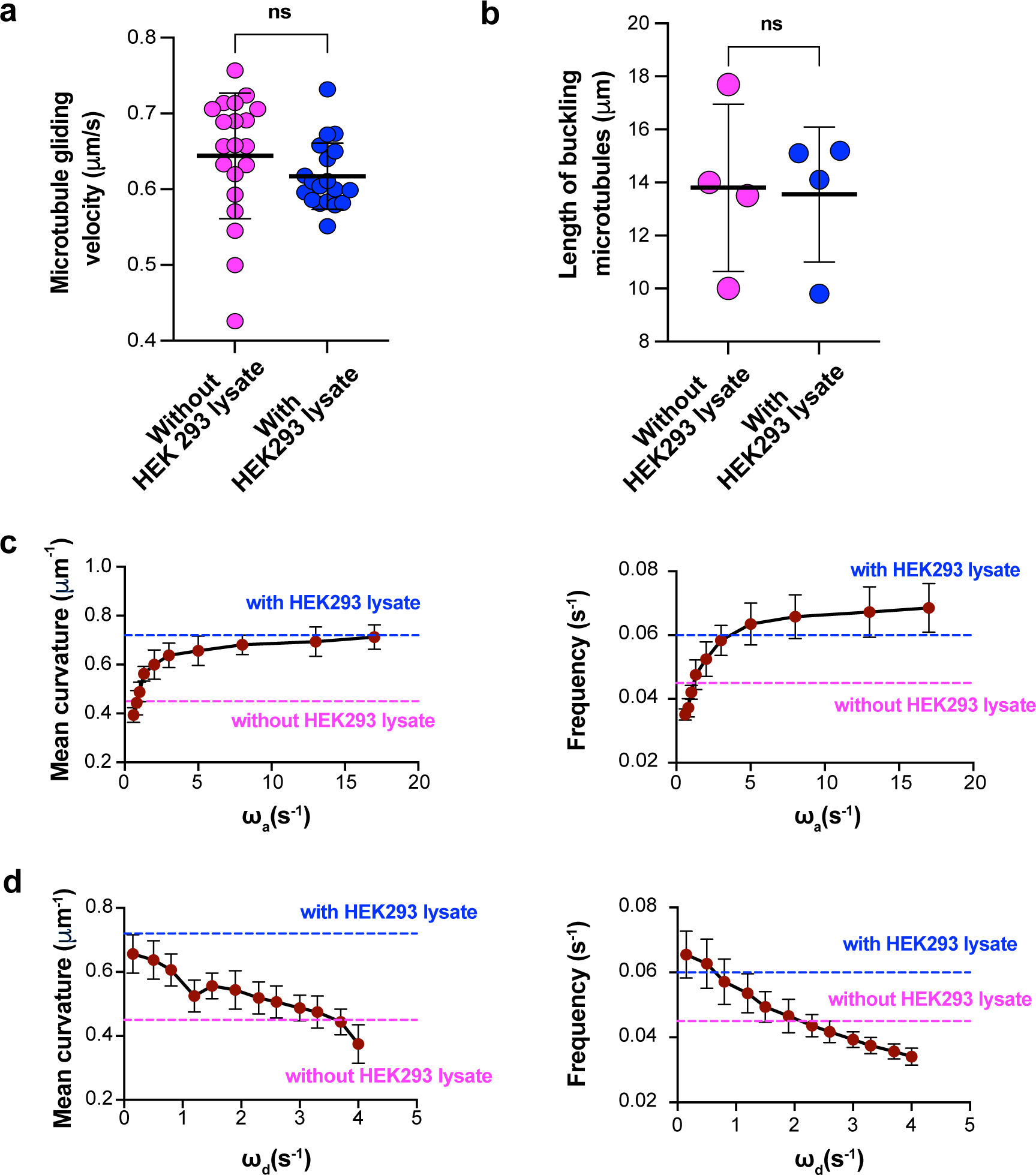
**a**, *Comparison of microtubule gliding velocity* with and without the addition of 20 µg ml^-1^ HEK293 cell lysate. Black lines represent mean and error bars represent the S.D. p= 0.2146 (not significant; ns) using unpaired t-test (n= 19 microtubules analyzed in each condition, from two independent experiments). **b**, *Length of buckling microtubules* compared for estimating mean curvatures of buckling microtubules with and without the addition of 20 µg ml^-1^ HEK293 cell lysate (Refer fig 7d). p= 0.9054 (not significant; ns) using unpaired t-test (n= 26 frames analyzed from 4 microtubules in each condition from three independent experiments). Mean curvature (left) and oscillation frequency (right) as a function of the motor attachment rate (ω_a_), in **7c** and motor detachment rate (ω_d_), in **7d** for a microtubule of L = 10 μm and L_p_ = 5 mm. The symbols and error bars indicate mean ± S.D respectively. Blue and pink dotted lines represent experimentally determined values with and without HEK293 lysate respectively.

