## Supplementary_Information for "Kinesin-induced buckling reveals the limits of microtubule self-repair"

**Affiliations:**
[1] Experimental Physics and Center for Biophysics, Saarland University, 66123 Saarbrücken, Germany
[2] Theoretical Physics and Center for Biophysics, Saarland University, 66123 Saarbrücken, Germany
[3] B CUBE - Center for Molecular Bioengineering, TUD Dresden University of Technology, 01307 Dresden, Germany
[4] Max Planck Institute of Molecular Cell Biology and Genetics, 01307 Dresden, Germany
[5] Cluster of Excellence Physics of Life, TUD Dresden University of Technology, 01062 Dresden, Germany
[6] PharmaScienceHub (PSH), 66123 Saarbrücken, Germany

**This PDF file includes:**
1. Supplementary text for simulation methods
2. Extended data table and supplementary references 3. Legends for supplementary movies

### Simulation Methods

**Description of the microtubule:**
Our numerical model describes the active bending of microtubules (MTs), which are represented as semi-flexible filaments subject to deformations from active forces generated by molecular motors. The energy of a given filament of length $L$ and bending rigidity $k_{\text{bend}} = L_p k_B T$ ($L_p$ being the persistence length of the filament) is calculated using the worm-like chain model [70, 71]:

$$E_{\text{filament}} = \frac{k_{\text{bend}}}{2} \int_0^L \left( \frac{\partial \theta(s)}{\partial s} \right)^2 ds, \tag{1}$$

where $s$ is the contour length along the filament, $\theta(s)$ is the local tangent angle, and $\partial\theta(s)/\partial s$ is the local curvature. To simulate filament dynamics numerically, the filament is discretized into $N$ segments of equal length ($\Delta L = \frac{L}{N} = 8\,\text{nm}$). The segments are assumed to be inextensible. The configuration of the filament is then parametrized by a set of angular coordinates $\theta_i$ corresponding to the orientation of each segment with respect to the $x$-axis. The discretized version of the Hamiltonian becomes:

$$E_{\text{filament}} = \frac{k_{\text{bend}}}{\Delta L} \sum_{i=1}^{N} \Big( 1 - \cos\left(\theta_i - \theta_{i-1}\right) \Big). \tag{2}$$

The angle of the first segment is fixed at $\theta_0 = 0$. For a straight filament aligned with the $x$-axis, $E_{\text{filament}} = 0$. The position of each node $\mathbf{x}_i$ is given by:

$$\mathbf{x}_i = \sum_{j=0}^{i} \Delta L \begin{pmatrix} \cos(\theta_j) \\ \sin(\theta_j) \end{pmatrix}. \tag{3}$$

**Active cross-linkers:**

We model the action of active molecular motors stochastically. Motors (modeled as cross-linkers) attach from a 2D grid underlying the filament. Each cross-linker on this grid can stochastically attach to a node on the filament. Once attached, the cross-linker applies a force like a linear spring:

$$E_{\text{crosslink}} = \frac{k_{\text{spring}}}{2} \sum_{i=1}^{n} ||\mathbf{x}_{\text{link},i} - \mathbf{x}_{\text{grid},i}||^2, \tag{4}$$

where $k_{\text{spring}}$ is the spring constant, $\mathbf{x}_{\text{link},i}$ is the node between adjacent segments of the filament, and $\mathbf{x}_{\text{grid},i}$ is the fixed point of the cross-linker on the grid. If the node is within a attachment radius $d_{\text{attach}}$, unbound motors can attach to the filament with rate $\omega_{\text{a}}$. While the attachment rate can be force-dependent in general (i.e. $\omega_{\text{a}}(F) = \omega_{\text{a,0}} \exp\left(-\frac{|F|}{f_{\text{a}}}\right)$ with $f_{\text{a}}$ being the characteristic attachment force), here we report the results for a constant $\omega_{\text{a}}$.

In contrast, the detachment rate depends on the force:

$$\omega_{\text{d}}(F) = \omega_{\text{d,0}} \exp\left(\frac{|F|}{f_{\text{d}}}\right), \tag{5}$$

where $f_{\mathsf{d}}$ is the characteristic detachment force and $|F|$ is the absolute value of the force exerted on the filament by the motor. To avoid unrealistically large forces and numerical instabilities, a maximum motor extension length of $40\,$nm was imposed. Once attached, motors can walk along the filament from the fixed end toward the free end. The step size $d_{\mathsf{h}}$ is constant ($d_{\mathsf{h}} = \Delta L = 8\,$nm). Similar to the detachment rate, the hopping rate is also force-dependent. If the force and motion direction are aligned the hopping rate is given by

$$\omega_{\mathsf{h}}(F) = \mathsf{min}\left[2\,\omega_{\mathsf{h,0}}, \omega_{\mathsf{h,0}}\left(1 + \frac{|F|}{f_{\mathsf{s}}}\right)\right], \tag{6}$$

otherwise

$$\omega_{\mathsf{h}}(F) = \mathsf{max}\left[0, \omega_{\mathsf{h,0}}\left(1 - \frac{|F|}{f_{\mathsf{s}}}\right)\right], \tag{7}$$

where $f_{\mathsf{s}}$ is the stall force and $\omega_{h,0}$ is the load-free hopping rate. In the simulation, we assume a separation of timescales: the relaxation of the filament is much faster than the timescale of motor dynamics. Therefore, after each motor update (attachment, detachment, or stepping), the shape of the filament is re-equilibrated. Nevertheless, this equilibrated configuration will change in the next time step since the status of motors evolves with time (including new attachments, new detachments, and motor position updates). The choice of the initial nodes to attach motors to the filament and their updates because of motor dynamics are carried out using the Gillespie algorithm. After each step, the energy of the system is minimized using a gradient descent method.

The mean force per motor is computed by evaluating the elastic stretching force of each motor, modeled as a linear spring. Specifically, for each active motor, we calculate the product of the spring constant and the displacement vector between the position of the motor on the filament and its fixed attachment point on the grid. The mean force per motor is then obtained by averaging these forces over all active motors bound to the filament. To determine the mean force per discrete filament segment, we first sum the force vectors from all motors that are attached to the same node of the filament. This vector sum represents the total force exerted on that segment. Finally, we compute the average over all such segments along the filament to obtain the mean force per discrete segment. To quantify the local curvature of the filament, we assign a curvature value to each internal node based on the positions of three successive nodes along the discretized filament. For each triplet of adjacent nodes, we compute the radius of the circle that passes through all three points. The local curvature at the central node is then defined as the inverse of this radius.

To investigate the spatial relationship between local curvature and force along the filament, we computed the cross-correlation function Corr$(\Delta x)$ between the local curvature $c(x,t)$ and the local force $f(x{\pm}\Delta x, t)$, evaluated at spatially offset positions along the filament. This function quantifies the correlation between curvature at position $x$ and force at a neighboring position $x{\pm}\Delta x$, yielding a value between 1 and -1, indicating positive and negative correlation, respectively.
The cross-correlation was calculated using the standard Pearson correlation coefficient formula: Corr$(\Delta x) = \frac{\langle (c(x,t)-\langle c\rangle)(f(x\pm\Delta x,t)-\langle f\rangle)\rangle}{\sigma_c \sigma_f}$. Here, the parameters enclosed within $\langle\rangle$ are obtained by averaging over filament segments at a given time $t$, and $\sigma_c$ and $\sigma_f$ are the standard deviations of the curvature and force, respectively.

#### Estimation of the steady-state number of bound motors:

To compute the steady-state number of motor proteins attached to a microtubule filament, we model motor binding and unbinding as a stochastic two-state Markov process influenced by an external force. Motors bind with a force-dependent attachment rate $\omega_{\mathrm{a}}(f) = \omega_{\mathrm{a},0} \exp\left(-\frac{|F|}{f_{\mathrm{a}}}\right)$ and detach with a rate $\omega_{\mathrm{d}}(f) = \omega_{\mathrm{d},0} \exp\left(\frac{|F|}{f_{\mathrm{d}}}\right)$ (see Eq. 5). $|F|$ is the absolute value of the average force exerted on the filament by the motors at different motor densities (refer Fig 6i). By analyzing the Markov chain balance of stochastic binding and unbinding events, the steady-state fraction of available motors which remain bound can be obtained as $\frac{1}{1+\frac{w_{d,0}}{w_{a,0}} \exp(|F|(1/f_d+1/f_a))}$. Given a surface motor density $\rho$, the number of motors geometrically available to interact with the filament is approximately $\sqrt{\rho}$, per micrometer.
Therefore, the steady-state number of motors that remain bound yields

$$N_{\text{steady}} = \frac{\sqrt{\rho}}{1 + \frac{w_{d,0}}{w_{a,0}} \exp(|F|(1/f_d + 1/f_a))}$$

In case of a constant (force-independent) $\omega_{\mathrm{a}}$, as in our simulations, the relation for the steady-state number of bound motors reduces to

$$N_{\text{steady}} = \frac{\sqrt{\rho}}{1 + \frac{w_{d,0}}{w_{a,0}} \exp(|F|/f_d)}$$

#### Estimation of forces acting on statically bent microtubules in experiments:

To estimate the force exerted on statically bent microtubules observed in experiments (Refer Fig 1c, right), we model the microtubule as a semiflexible filament confined to two dimensions and clamped at one end [41]. The full shape of the microtubule is extracted from microscopy images. An external force with unknown $x$- and $y$-components is assumed to act at the free end. Our goal is to determine the force that best reproduces the observed static configuration under these boundary conditions. For the numerical simulations, the filament is assumed to have a given bending rigidity and is discretized into segments, as described above. To estimate the applied force, we fix the position and orientation (tangent angle) at the fixed end, the components of a trial force at the free end and obtain the equilibrium configuration of the filament. If the applied trial force differs from the true value, the resulting shape evolves away from the experimentally observed configuration and the position of the free end changes. Thus, for each trial force we compute the Euclidean distance between the simulated and observed position of the free end. By scanning across a range of trial force values, we identify the one that minimizes this distance as the optimal force reproducing the observed filament shape under the given constraints.

#### Extended data table and supplementary references:

[70] A. Marantan, L. Mahadevan, ”Mechanics and statistics of the worm-like chain”, Am. J. Phys. 86, 2018.

[71] I. Weber, C. Appert-Rolland, G. Schehr and L. Santen, ”Non-equilibrium fluctuations of a semi-flexible filament driven by active cross-linkers”, EPL 120, 38006, 2017.

#### Legends for supplementary movies:

Supplementary video 1: Different microtubule bending events in live PtK2 cells. Videos showing dynamic buckling, microtubules persisting in the bent shape, loop formation and breakage in Ptk2 cells (endogenous tubulin-eGFP tag; represented here in magenta). Movies are at 10 fps.

Supplementary video 2: Different microtubule bending events recaptured in our dynamic buckling assay. Using our in vitro assay, we recapture microtubule behaviours as seen in cells- regular flagella-like oscillations, loop formation, beating as well as pivoting. Movies are at 20 fps.

Supplementary video 3: Trace of mean curvature over time (left) of a microtubule (right) as it buckles. Movie is at 20 fps.

Supplementary video 4: Gliding microtubule with incorporation. Movie is at 5 fps.

Supplementary video 5: Buckling microtubule with incorporation. Movie is at 5 fps.

Supplementary video 6: Looping microtubule with incorporation. Movie is at 20 fps.

Supplementary video 7: Buckling microtubules in vitro breaking in the presence of 5 µM unlabeled free tubulin. Movie are at 20 fps.

Supplementary video 8: Compilation of buckling and looping microtubules breaking at points of high curvature. Movies are at 20 fps.

Supplementary video 9: Evolution of microtubule shape in the buckling regime during simulations. The motor density is set to ρ= 278 per µm2, with all other parameters at their default values as listed in ED. Table 1.

Supplementary video 10: Evolution of microtubule shape in the loop and knot formation regime during simulations. The motor density is set to ρ= 494 per µm2, with all other parameters at their default values as listed in ED.Table 1.

Supplementary video 11: Evolution of microtubule shape in the beating regime during simulation. The motor density is set to ρ= 123 per µm2, with all other parameters at their default values as listed in ED.Table 1.

Supplementary video 12: Timelapse of double-anchored microtubule subjected to pulling action of motors in a double-anchored assay. Movie is at 20 fps.

Supplementary video 13: Comparison of buckling microtubule- in the absence of (left).vs in the presence of 20 µg/ml HEK293 wild type lysate. Movies are at 20 fps.
